# Incremental language comprehension difficulty predicts activity in the language network but not the multiple demand network

**DOI:** 10.1101/2020.04.15.043844

**Authors:** Leila Wehbe, Idan Asher Blank, Cory Shain, Richard Futrell, Roger Levy, Titus von der Malsburg, Nathaniel Smith, Edward Gibson, Evelina Fedorenko

## Abstract

What role do domain-general executive functions play in human language comprehension? To address this question, we examine the relationship between behavioral measures of comprehension and neural activity in the domain-general “multiple demand” (MD) network, which has been linked to constructs like attention, working memory, inhibitory control, and selection, and implicated in diverse goal-directed behaviors. Specifically, fMRI data collected during naturalistic story listening are compared to theory-neutral measures of online comprehension difficulty and incremental processing load (reading times and eye-fixation durations). Critically, to ensure that variance in these measures is driven by features of the linguistic stimulus rather than reflecting participant-or trial-level variability, the neuroimaging and behavioral datasets were collected in non-overlapping samples. We find no behavioral-neural link in functionally localized MD regions; instead, this link is found in the domain-specific, fronto-temporal “core language network”, in both left hemispheric areas and their right hemispheric homologues. These results argue against strong involvement of domain-general executive circuits in language comprehension.

## Introduction

Human language comprehension encompasses a host of complex computations, from perceptual (auditory, visual or, in the case of Braille, haptic) processing, to word recognition, to recovering the semantic and syntactic dependency structures linking words together, to constructing discourse-level representations, and making pragmatic inferences. A major goal of both behavioral psycholinguistics and cognitive neuroscience of language is to understand which cognitive mechanisms support language comprehension, and whether and how these mechanisms are shared with other (non-linguistic) cognitive functions.

Psycholinguists have long invoked *domain-general constructs* when discussing lexical access and syntactic/semantic dependency formation, from storage and retrieval of information from working memory, to updating focal attention, inhibiting irrelevant information, selecting an option among alternatives, and predictive processing (Abney & Johnson, 1991; Fedorenko, Gibson, & Rohde, 2006, 2007; Gernsbacher, 1993; E. Gibson, 1998, 2000; Gordon, Hendrick, & Levine, 2002; Johnson-Laird, 1983; King & Just, 1991; Lewis & Vasishth, 2005; Lewis, Vasishth, & Van Dyke, 2006; McElree, 2000, 2001; Novick, Kan, Trueswell, & Thompson-Schill, 2009; Rasmussen & Schuler, 2018; Resnik, 1992; Rodd, Johnsrude, & Davis, 2010; Schuler, AbdelRahman, Miller, & Schwartz, 2010; van Schijndel, Exley, & Schuler, 2013; Vergauwe, Barrouillet, & Camos, 2010; Waters & Caplan, 1996, *inter alia*). If some linguistic processes require these or other domain-general operations, does it mean that language shares neural mechanisms with other domains?

It has long been known that language processing recruits particular neural circuitry (Broca, 1861; Geschwind, 1970; Wernicke, 1874). However, prior cognitive neuroscience work has argued both (1) that some of this circuitry (e.g., “Broca’s area”) may not be specialized for language processing *per se*, but rather used for broader cognitive functions—like hierarchical syntactic structure building—that operate not only in language but also in other domains like music, mathematics, and action planning (Anderson, 2010; Fadiga, Craighero, & D’Ausilio, 2009; Fitch & Martins, 2014; Friedrich & Friederici, 2009; Patel, 2003, 2012; Rodriguez & Granger, 2016; Slevc, Rosenberg, & Patel, 2009; Tettamanti & Weniger, 2006; *inter alia*, see Fedorenko & Blank, 2020 for a review); and (2) that language processing relies on a more spatially distributed network, extending beyond the “classic” language areas, that includes regions traditionally associated with domain-general executive control (Mesulam, 1998; Kaan and Swaab, 2002; Kuperberg et al., 2003; Novick et al., 2005; Rodd et al., 2005a; Thompson-Schill et al., 2005; Novais-Santos et al., 2007; January et al., 2009b; Peelle et al., 2010; Rogalsky and Hickok, 2011; Wild et al., 2012; McMillan et al., 2012, 2013; Nieuwland et al., 2012; Blumstein and Amso, 2013; Hsu and Novick, 2016; *inter alia*). Hypotheses from psycholinguistics, cognitive science, and cognitive neuroscience therefore converge to predict a role for domain-general executive resources in human language comprehension.

Within the human brain, the most plausible place to look for domain-general recruitment is in the fronto-parietal / cingulo-opercular “multiple demand (MD)” network, which supports a broad range of executive functions, including inhibitory control, attentional selection, conflict resolution, and maintenance and manipulation of task sets (Duncan, 2010; Fedorenko, Duncan, & Kanwisher, 2013). Indeed, MD regions have been shown to be sensitive to linguistic processing difficulty (e.g., due to ambiguity or complexity) across diverse manipulations (Kuperberg et al., 2003; Rodd et al., 2005a; Novais-Santos et al., 2007; January et al., 2009b; Peelle et al., 2010; Nieuwland et al., 2012; McMillan et al., 2013 *inter alia*). Further, activity in this network has been shown to correlate positively with reaction times—a behavioral measure of processing difficulty—across tasks (Taylor, Rastle, & Davis, 2014; Yarkoni, Barch, Gray, Conturo, & Braver, 2009). If indeed MD regions register processing load during language comprehension, this would support the hypothesis that domain-general resources are engaged in language comprehension.

The ability of prior work to bear on this hypothesis is limited by two factors. First, language comprehension effort has typically been studied by relating theory-driven linguistic variables (e.g., word frequency, word predictability, structural complexity, constituent length, etc.) to neural activity (Mazoyer et al., 1993; Stowe et al., 1998; Vandenberghe et al., 2002; Friederici et al., 2003; Dronkers et al., 2004; Humphries et al., 2006; Brennan et al., 2010, 2016; Pallier et al., 2011; Rogalsky and Hickok, 2011; Brennan and Pylkkänen, 2012; Willems et al., 2016; Henderson et al., 2016; Lopopolo et al., 2017; Nelson et al., 2017; *inter alia*). Despite the critical role of theory in understanding human cognition, theory-driven variables are only as good as the underlying theory and can only be expected to capture a fraction of the language comprehension effort given the multi-faceted nature of language. Such variables may fail to characterize some components of language comprehension and thereby underestimate the extent to which some neural circuits are implicated in comprehension. Second, prior work, including many of the aforementioned studies purportedly showing MD involvement in language comprehension, has generally relied on language stimuli cleverly constructed to directly manipulate some aspect of language processing difficulty and has often included explicit tasks on top of language comprehension, like making judgments about sentences or deciding whether a sentence matches a picture (e.g. Friederici et al., 2003; Fiebach et al., 2004; Rodd et al., 2005a; Bilenko et al., 2008; Kuperberg et al., 2008; Snijders et al., 2009; Blank et al., 2016). Such hand-constructed stimuli and tasks are very different from natural comprehension “in the wild”, and may inadvertently trigger recruitment of domain-general problem solving and task strategizing mechanisms due to their artificial nature and extraneous task demands (Campbell & Tyler, 2018; Diachek, Blank, Siegelman, Affourtit, & Fedorenko, in press; Hasson, Egidi, Marelli, & Willems, 2018; Hasson & Honey, 2012). Such stimuli and tasks might thus overestimate MD involvement in language comprehension, especially given the sensitivity of MD regions to task demands (D’Esposito & Postle, 2015; Miller & Cohen, 2001; Sreenivasan, Curtis, & D’Esposito, 2014). MD recruitment for language processing would therefore be better supported if an MD response to theory-neutral measures of comprehension difficulty could be shown under more naturalistic experimental conditions.

Therefore, in this study, to test the hypothesis of domain-general executive involvement in language comprehension, we use context-rich, naturalistic language stimuli presented without any extraneous tasks and correlate (1) experimentally-obtained behavioral reaction time measures of language processing difficulty during reading, with (2) fMRI measures of activity in the domain-general MD network. To increase the interpretability of such correlations, we compare them to brain-behavior correlations based on a different functional network: the domain-specific, fronto-temporal “core language network”. This network serves as a good comparison for the MD network because it robustly engages in comprehension (during both listening and reading) but shows little to no engagement in other high-level cognitive processes (Binder, 1997; Deniz, Nunez-Elizalde, Huth, & Gallant, 2019; Fedorenko, Behr, & Kanwisher, 2011; Fedorenko & Blank, 2020; Fedorenko, Hsieh, Nieto-Castanon, Whitfield-Gabrieli, & Kanwisher, 2010; Fedorenko & Varley, 2016; Jung-Beeman, 2005). Below, we describe and justify the main design features of our experiment.

Our use of behavioral reading data as a global proxy for comprehension difficulty follows a standard psycholinguistic paradigm that investigates how reaction times vary in response to linguistic materials whose comprehension requires different kinds of (hypothesized) computations, in either experimentally constructed materials (e.g., Frazier and Rayner, 1987; Clifton and Frazier, 1989; Gibson, 1991, 1998; Grodner et al., 2002; Levy, 2008), or naturalistic ones (e.g. Demberg and Keller, 2008; Smith and Levy, 2013). Although incremental reading data are known to have a complex relationship to mental states (Posner, 1980, 2016; Remington, 1980; Klein and Farrell, 1989; Wright and Ward, 2008; *inter alia*) and be sensitive to non-linguistic factors like general attention, sensory/perceptual processing, motor control, and task-related strategizing (Kaakinen & Hyönä, 2010; Kennedy, 2000; Rayner, 1998; Schotter, Tran, & Rayner, 2014), a premise underlying most psycholinguistic work in this domain is that incremental behavioral measures of reading effort track language-related comprehension difficulty with sufficient reliability such that they can be used to validate theories of human sentence comprehension (M. A. Just & Carpenter, 1980; Lewis et al., 2006; Mitchell, 1984; Rayner, 1977, 1978, 1998). Furthermore, our experimental design reduces the influence of idiosyncratic processes such as attention fluctuations by (1) aggregating reading data from many participants; (2) separating the samples that provide behavioral data from the sample providing the neuroimaging data; and (3) using different presentation modalities across the behavioral (visual) and fMRI (auditory) paradigms (cf. Henderson et al., 2015). This design is intended to distill stimulus-related, generalizable variation in comprehension difficulty: because attention, sensory/perceptual, motor, and task variables are unlikely to co-vary between participants and presentation modalities, any correlation between behavioral and neural measures in this design is most plausibly due to the linguistic content of the stimuli themselves, and thus is most plausibly driven by language comprehension effort.

We consider two different behavioral responses—self-paced reading (SPR, Aaronson and Scarborough, 1977; Just et al., 1982) and eye-tracking during reading (ET, Rayner, 1998), from two large, existing datasets (Futrell et al., 2018 and von der Malsburg et al., unpublished). These measures of comprehension effort serve as theory-neutral, broad-coverage estimates of computational load during language comprehension, since they should permit detection of any mechanisms that contribute to processing latencies, even if their role is not yet captured by any existing theory.

When correlating these measures with neuroimaging data, we consider the detailed time-course of activation during listening, rather than an aggregate measure averaging across the entire stimulus, or parts of the stimulus. The time-varying fMRI data enable us to exploit relatively fine-grained variation in incremental processing difficulty that may be attenuated in aggregate measures. In addition, we infer the hemodynamic response from the data, in order to address individual and regional variation in the underlying hemodynamic response (Handwerker, Ollinger, & D’Esposito, 2004). Finally, we employ non-parametric hypothesis tests on out-of-sample data, in order to increase the statistical robustness of the results and reduce the risk of replication failure (Eklund, Andersson, Josephson, Johannesson, & Knutsson, 2012; Menke & Martinez, 2004).

To foreshadow our results, whereas we find that reading latencies predict neural activity in the core language network, we do not find that reading latencies predict neural activity in the MD network. This finding supports the hypothesis that incremental processing effort during naturalistic language comprehension is largely restricted to neural circuits (and, by extension, cognitive resources) that are specialized for language comprehension, with little role played by domain-general executive systems.

## Materials and methods

### Short stories

We use the Natural Stories Corpus (Futrell et al., 2018; data downloaded from https://github.com/languageMIT/naturalstories.git), which contains ten stories that were constructed from existing, publicly available texts (fairy tales, short stories, and Wikipedia articles) but edited so as to make comprehension difficulty more variable than in fully natural texts. The dataset includes recordings of these stories by two native English speakers (one male, E.G., and one female).

### Behavioral self-paced reading data

The Natural Stories Corpus includes self-paced reading data from 181 native English speaking participants recruited through Amazon.com’s Mechanical Turk. Participants gave informed consent in accordance with the Internal Review Board at the Massachusetts Institute of Technology (MIT) and were paid for their participation. Participants read stories in a moving-window self-paced word-by-word reading paradigm, where a button has to be pressed to reveal each subsequent word. The time spent on each word provides an overall estimate of processing difficulty at that point in the sentence/story. Each story was followed by 6 multiple-choice comprehension questions and if a participant answered fewer than 5 questions correctly, their reading time data for that story were excluded. Outlier reading times of less than 100ms or more than 3,000ms were also excluded. These exclusion criteria were the ones followed by Futrell et al. (2018). Reading times were aggregated across participants for each word. As a result, for each word in each story, we have a single (average) reading time.

### Behavioral eye-tracking study

Forty native English speaking participants recruited from the University of California, San Diego (UCSD) undergraduate population gave informed consent in accordance with the Internal Review Board at UCSD and were paid for their participation. They read the stories in an eye-tracking paradigm. A tower-mounted EyeLink 1000 eye-tracker recorded eye movements as participants read the stories presented a few sentences at a time (the boundaries among the story fragments and lines within fragments differed across participants so as to vary the words that span the screen-change and line boundaries). Each story was followed by two true/false comprehension questions. Software for automatic correction of eye fixations was used to repair data recorded with imperfect eye-tracker calibration (A. L. Cohen, 2013). A set of heuristics were used to detect and remove episodes of track loss, poor-quality data, and episodes where reader merely skimmed the text. In particular, fixations were removed when 1) the previous and/or subsequent fixations were five or more words away which is indicative of skimming (all the skipped words were also removed from the subject’s data in this case), 2) initial fixations on a new page of text occurred on words that were not at the beginning of the text, 3) the fixations could not be mapped to any word, or 4) consecutive fixations were moved in different directions by Cohen correction (J. Cohen, Cohen, West, & Aiken, 2013).^1^ For each word, four canonical eye-tracking measures were calculated (first pass regression, regression path duration, first pass reading time, and first fixation progressive) which are believed to index different perceptual and linguistic processes involved in reading, ranging from word recognition to high-level discourse integration (Rayner, 1998; Vasishth et al., 2013). Eye-tracking measures were aggregated across participants for each word. As a result, for each word in each story, we have four (average) eye-tracking measures.

### fMRI experiment

#### Participants

42 right-handed native English speakers (average age 22.7, SD = 3.3; 24 females) from the MIT community gave informed consent in accordance with the Internal Review Board at MIT and were paid for their participation. (Subsets of this dataset were used by Blank & Fedorenko (2017), Blank, Kanwisher, & Fedorenko (2014) and Shain, Blank, van Schijndel, Schuler, & Fedorenko, 2020).

#### General approach

Each participant listened to a subset of the stories from Futrell et al. (2018) and performed one or more “localizer” tasks (e.g. Saxe et al., 2006) used to identify the brain networks of interest.

#### Critical task

Participants listened to the recordings of the spoken stories. Each story corresponded to one fMRI run. Eight of the ten stories were used, and any given participant heard between 2 and 8 stories (average=4; two stories: n=12, three stories: n=13, four stories: n=2, five stories: n=4, six stories: n=5, seven stories: n=1, eight stories, n=5). Each story lasted between 4.5 and 6 min. Participants were asked to listen attentively. At the end of each story, a set of six two-alternative forced-choice comprehension questions appeared one by one, and participants answered by pressing one of two buttons. These questions were designed to be challenging and required attentive listening and the ability to respond quickly. On average, participants failed to provide an answer to 11.5% of the questions (SD = 15.2%) and, on the remaining questions, their mean accuracy was 83.5% (SD = 10.1%). (Comprehension data were available for 33 participants: they were lost for 2 participants, not recorded for 3 participants due to a script error, and not collected for 4 participants who listened to the stories as part of a larger experiment for which the design did not include comprehension questions). A binomial test for each participant (uncorrected across participants) showed that all but one participant demonstrated above-chance accuracy (*p* < 0.01). (In the supplementary materials, we report our main analysis restricted to participants with very good performance, which revealed the same general pattern of results (compare **Figure 3b** and **Supplementary Figure 3**.)

**Figure 1:**
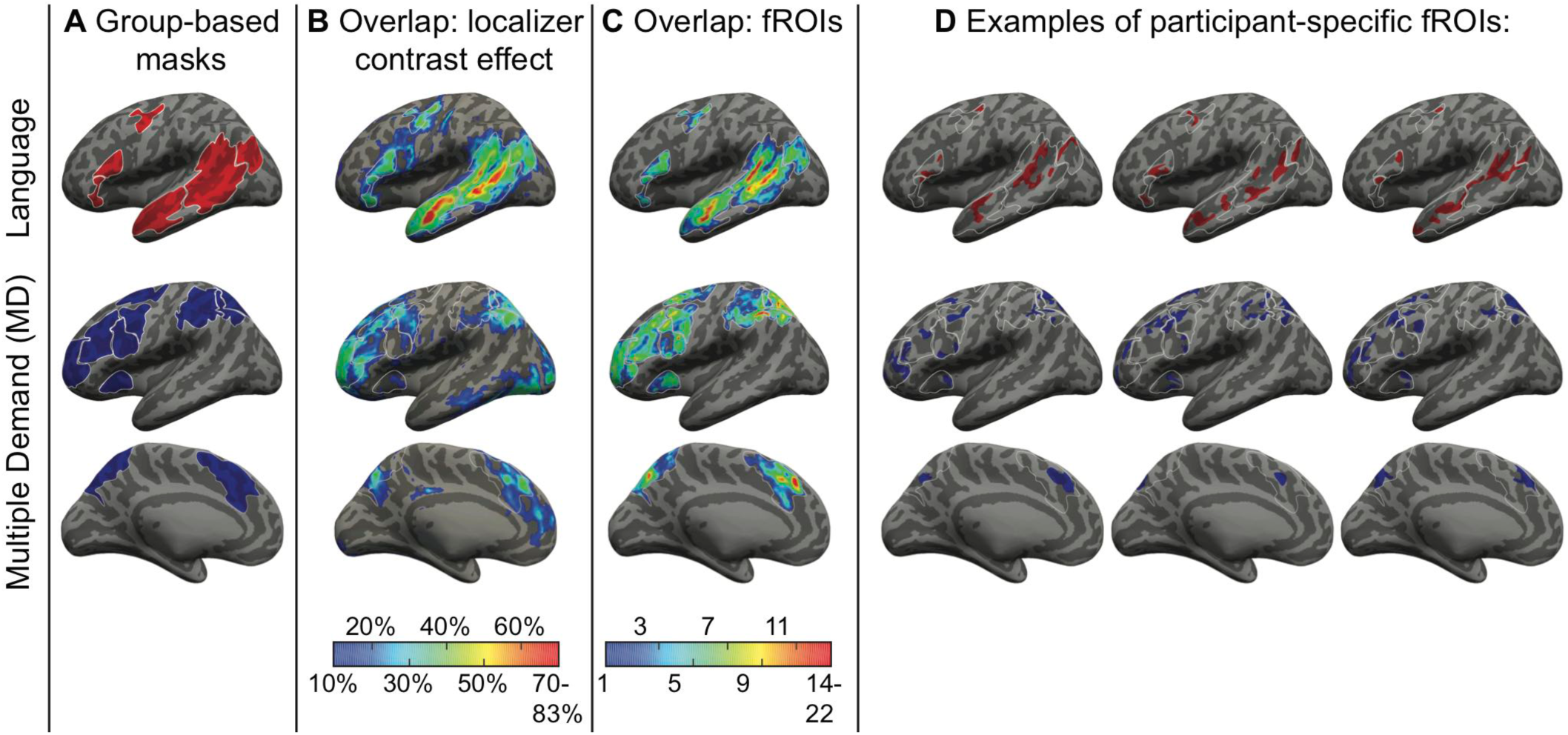
Defining participant-specific fROIs in the language (top) and MD (bottom) networks (only the left-hemisphere is shown). All images show approximated projections from functional volumes onto the surface of an inflated brain in common space. (A) Group-based masks used to constrain the location of fROIs. Contours of these masks are depicted in white on all brains in (B)-(D). (B) Overlap maps of localizer contrast effects (Sentence > Nonwords for the language network, Nonwords > Sentences for the MD network) across the 42 participants in the current sample (these maps were not used in the process of defining fROIs and are shown for illustration purposes). Each non gray-scale coordinate is colored according to the percentage of participants for whom that coordinate was among the top 10% of voxels showing the strongest localizer contrast effects across the nerocortical gray matter. (C) Overlap map of fROI locations. Each non gray-scale coordinate is colored according to the number of participants for whom that coordinate was included within their individual fROIs. (D) Example fROIs of three participants. Apparent overlap across language and MD fROIs within an individual is illusory and due to projection onto the cortical surface. Note that, because data were analyzed in volume (not surface) form, some parts of a given fROI that appear discontinuous in the figure (e.g., separated by a sulcus) are contiguous in volumetric space.

**Figure 2:**
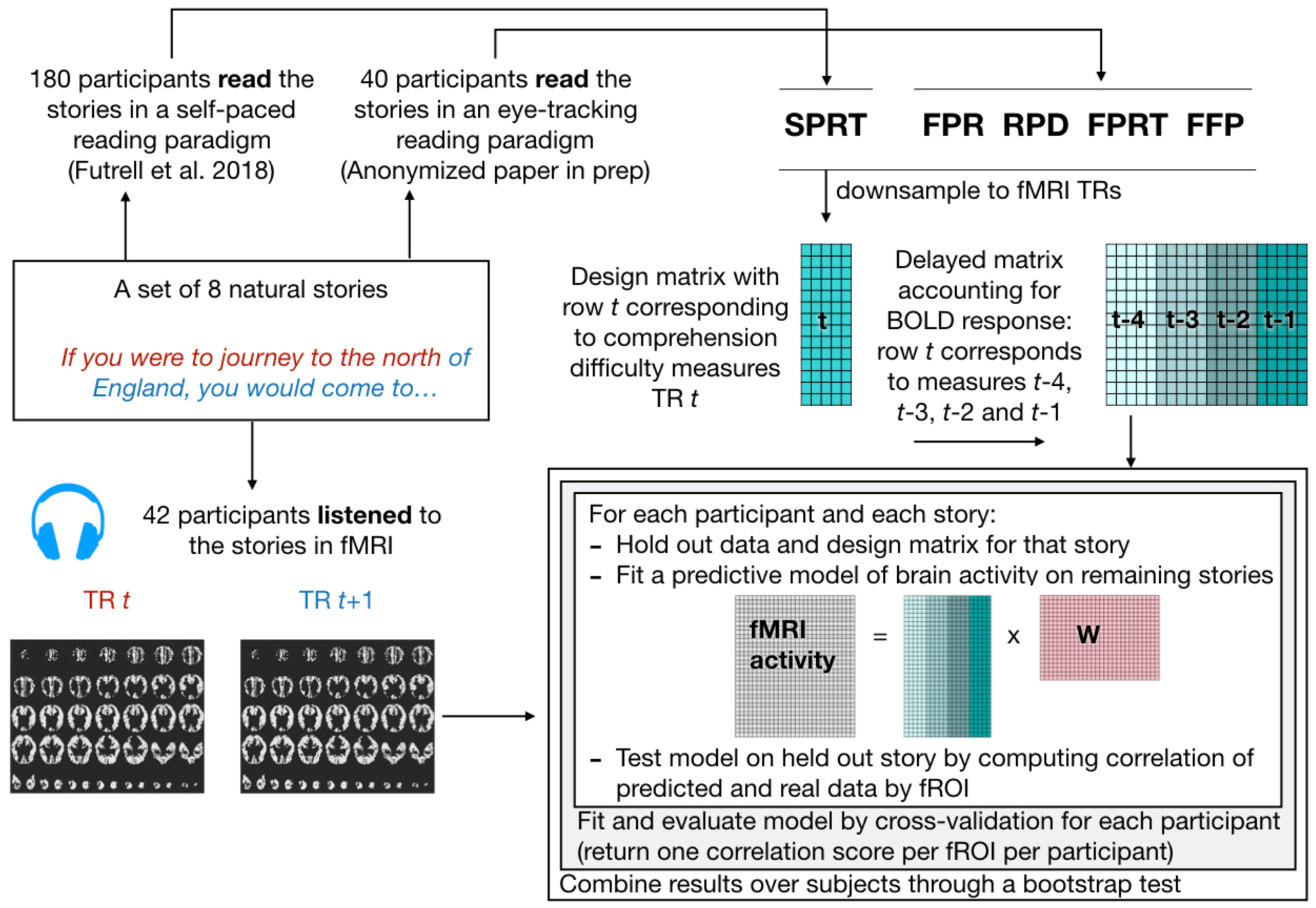
Diagram of approach detailing the combination of data from three experiments (fMRI, self-paced reading, and eye-tracking) with an encoding model approach. Comprehension difficulty measures are subsampled to the timing of the fMRI TRs and delayed to account for the BOLD response. For each subject, a cross-validation procedure is used where a story is held-out and a predictive model of brain activity as a linear combination of the comprehension difficulty measures is learned. The model is tested on the held-out story. Correlation of predicted and real data is computed for the held-out story; these value are then averaged across all cross-validation folds, resulting in an average correlation by subject and fROI (as well as fROI group). The cross-subjects results are combined using a bootstrap test.

**Figure 3:**
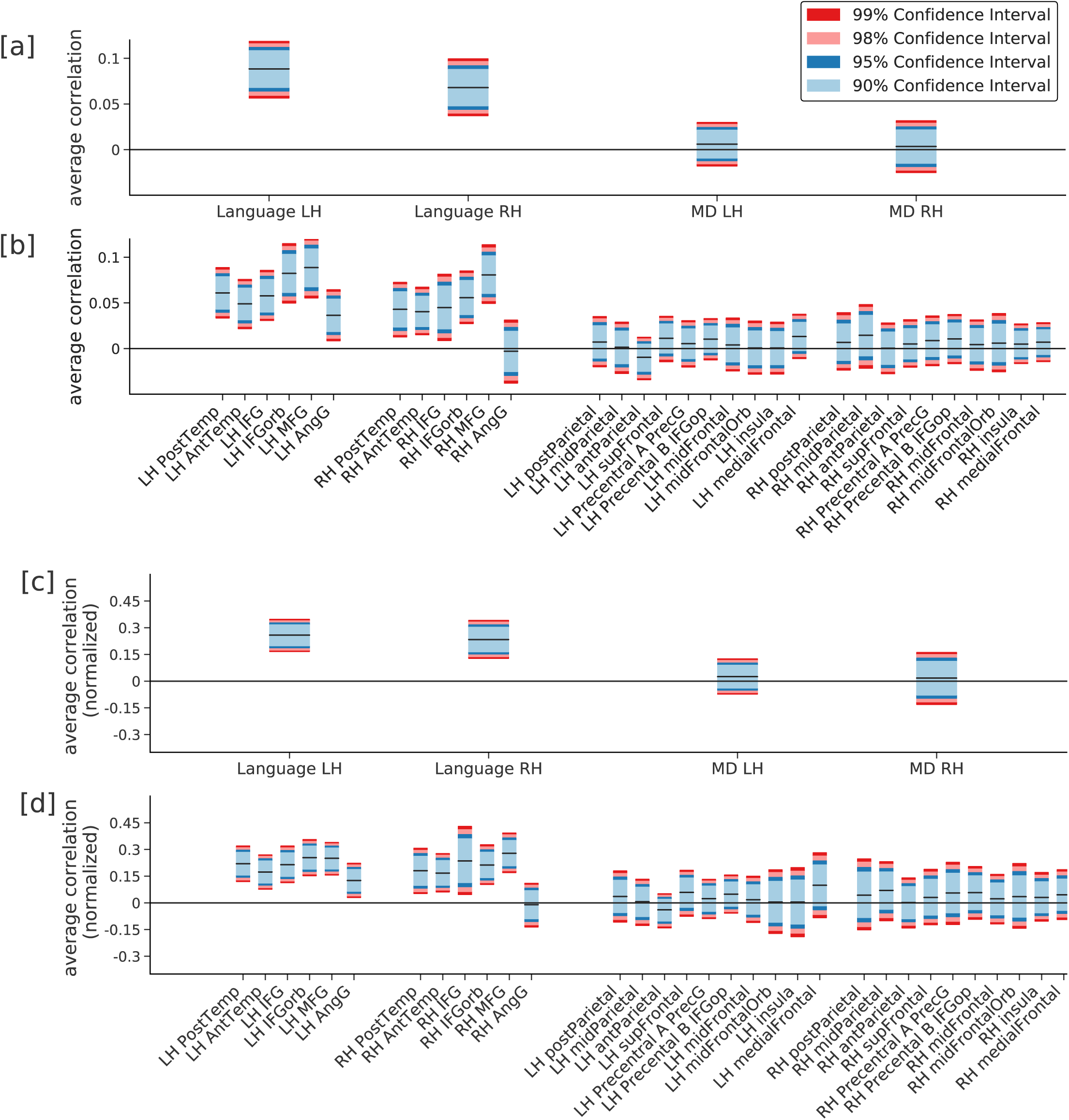
Average (unnormalized and normalized) correlation between activity predicted as a function of comprehension difficulty (estimated using a combination of self-paced reading times and eye-tracking measures) and real held-out activity, normalized by the estimated reliability of the signal for each fROI group ([a] unnormalized and [c] normalized) and each fROI ([b] unnormalized and [d] normalized). Performance was averaged across the 42 participants and bootstrap confidence intervals were constructed. Reading times predict the activity in left and right language fROIs, but not in MD fROIs.

#### Localizer tasks

All participants also performed an independent localizer task. This task was used to functionally identify the two networks of interest: the MD network, and the language network. We use the task described in detail in Fedorenko et al. (2010). Briefly, we used a reading task that contrasted sentences and lists of unconnected, pronounceable nonwords in a standard blocked design with a counterbalanced order across runs. Stimuli were presented one word / nonword at a time (for timing parameters, see **Table 1**). Eighteen participants read the materials passively (a button-press task at the end of each trial was included in order to maintain alertness); for the remaining 24 participants, each trial ended with a memory probe, i.e., a word / nonword, and they had to indicate (via a button press) whether or not this probe had appeared in the preceding sentence / nonword sequence. Each participant completed 2-4 runs of the localizer task. (A version of this localizer is available from https://evlab.mit.edu/funcloc/download-paradigms.) Because this localizer was originally designed to identify the core language network, we begin by describing how it was used to localize this network; we turn to the MD network next.

**Table 1:** Summary of the procedural and timing details for the different versions of the language localizer used in the current study.

|  | Version |  |  |  |
| --- | --- | --- | --- | --- |
|  | I | II | III | IV |
| Number of participants | 24 | 7 | 6 | 5 |
| Task: passive reading / memory probe? | PR | MP | MP | MP |
| Conditions | Sentences,<br>Nonwords | Sentences,<br>Word lists,<br>Nonwords | Sentences,<br>Nonwords | Sentences,<br>Word lists,<br>Nonwords |
| Words / nonwords per trial | 12 | 12 | 12 | 8 |
| Trial duration (ms) | 6000 | 6000 | 6000 | 4800 |
| Fixation | 100 | 300 | 300 | 300 |
| Presentation of each word / nonword | 450 | 350 | 350 | 350 |
| Memory probe | --- | 1000 | 1000 | 1350 |
| Fixation | 500 | 500 | 500 | 350 |
| Trials per block | 3 | 3 | 3 | 5 |
| Block duration (s) | 18 | 18 | 18 | 24 |
| Blocks per condition per run | 8 | 6 | 8 | 4 |
| Fixation block duration (s) | 14 | 18 | 18 | 16 |
| Number of fixation blocks per run | 5 | 4 | 5 | 3 |
| Total run time (s) | 358 | 396 | 378 | 336 |
| Number of runs | 2 | 2-3 | 2 | 3-4 |

The Sentences > Nonwords localizer contrast targets brain regions that support high-level language comprehension. This contrast generalizes across tasks (Fedorenko et al., 2010; Scott, Gallée, & Fedorenko, 2017) and presentation modalities (reading vs. listening; e.g., Fedorenko et al., 2010; Braze et al., 2011; Vagharchakian et al., 2012; Scott et al., 2017; Deniz et al., 2019).

All the regions identified by this contrast show sensitivity to lexico-semantic processing (e.g., stronger responses to real words than nonwords) and combinatorial syntactic and semantic processing (e.g., stronger responses to sentences and Jabberwocky sentences than to unstructured word and nonword sequences) (Bautista & Wilson, 2016; Blank et al., 2016; Fedorenko et al., 2010; E Fedorenko, Nieto-Castanon, & Kanwisher, 2012; Fedorenko, Blank, Siegelman, & Mineroff, 2020; Fedorenko et al., 2016; Heim, Eickhoff, & Amunts, 2008; Keller, Carpenter, & Just, 2001; Mineroff, Blank, Mahowald, & Fedorenko, 2018; Mollica et al., 2020; Rodd et al., 2005a). The Sentences > Nonwords contrast encompasses all of these processes, but narrower contrasts that target a subset of them identify the same cortical network (e.g. Fedorenko et al., 2010), suggesting that all the regions in the fronto-temporal language network support all of these high-level linguistic processes (for discussion, see Fedorenko, (in press) and Fedorenko, Mineroff, Siegelman, & Blank, (2018). In addition, the same network is identified by broader contrasts that do not subtract out phonological processing and also include pragmatic and discourse-level processes (e.g., a contrast between natural spoken paragraphs and their acoustically degraded versions or paragraphs in an unfamiliar language; (Ayyash, D.*, Malik Moraleda, Galleé, J., Z., Jouravlev, & Fedorenko, in prep.; Scott et al., 2017)). Finally, this localizer also identifies right-hemisphere homologues of the classic, left-hemisphere language regions (e.g., Mahowald and Fedorenko, 2016), which we included here because our other network of interest (the MD network) is bilateral and because right-hemisphere language regions have been previously implicated in several aspects of language comprehension (Deniz et al., 2019; Huth et al., 2016; Jung-Beeman, 2005; Wehbe et al., 2014).

To identify MD regions, we used the reverse, Nonwords > Sentences, contrast, which targets regions that increase their response during the more effortful reading of nonwords compared to that of sentences. This “cognitive effort” contrast robustly engages the MD network, can reliably localize it, and generalizes across a wide array of stimuli and tasks, both linguistic and non-linguistic (Fedorenko et al., 2013; Mineroff et al., 2018). We verified that the MD regions thus localized robustly respond to a difficulty (memory load) manipulation in a non-linguistic, visuo-spatial working-memory task, for a subset of 36 participants for whom data for this task had been collected: all regions showed a stronger response to a harder condition than to an easier condition (dependent samples *t*(35)>3.84, *p*<10^−6^, false discovery rate corrected for the number of regions; Cohen’s *d*>0.30, computed based on an independent samples formula, see **Supplementary Figure 1**). (In the supplementary materials, we additionally report our main analysis restricted to the 36 participants for whom the visuo-spatial working memory task data had been collected using the Hard > Easy contrast in that task to localize the MD regions. This analysis revealed the same pattern of results as in the main analysis where the Nonwords > Sentences contrast was used (compare **Figure 3b** and **Supplementary Figure 2**).)

### fMRI data acquisition

Structural and functional data were collected on the whole-body 3-Tesla Siemens Trio scanner with a 32-channel head coil at the Athinoula A. Martinos Imaging Center at the McGovern Institute for Brain Research at MIT. T1-weighted structural images were collected in 176 sagittal slices [1 mm isotropic voxels; repetition time (TR): 2,530 ms; echo time (TE): 3.48 ms]. Functional BOLD data were acquired using an echo planar imaging sequence with a flip angle of 90° and applying generalized autocalibrating partially parallel acquisition with an acceleration factor of two. Images were collected in 31 near-axial slices, acquired in an interleaved order with a 10% distance factor [in-plane resolution: 2.1×2.1 mm; slice thickness: 4 mm; field of view: 200 mm in the phase encoding anterior to posterior (A >> P) direction; matrix size: 96×96; TR: 2,000 ms; TE: 30 ms]. Prospective acquisition correction (Thesen, Heid, Mueller, & Schad, 2000) was used to adjust the positions of the gradients based on the subject’s head motion one TR back. The first 10 s of each run was excluded to allow for steady-state magnetization.

### fMRI data preprocessing

#### Spatial preprocessing

Data preprocessing was carried out with SPM5 and custom MATLAB scripts. (Note that SPM was only used for preprocessing and basic first-level modeling of the localizer data, aspects that have not changed much in later versions; we used an older version of SPM because data for this study are used across other projects spanning many years and hundreds of participants, and we wanted to keep the SPM version the same across all the participants.) Preprocessing of anatomical data included normalization into a common space (Montreal Neurological Institute (MNI) template, resampling into 2 mm isotropic voxels, and segmentation into probabilistic maps of the gray matter, white matter (WM) and cerebrospinal fluid (CSF). Preprocessing of functional data included motion correction, normalization, resampling into 2 mm isotropic voxels, smoothing with a 4 mm FWHM Gaussian kernel and high-pass filtering at 200s.

#### Temporal preprocessing

Additional preprocessing of data from the resting state and story comprehension runs was carried out using the CONN toolbox (Whitfield-Gabrieli & Nieto-Castanon, 2012) with default parameters, unless specified otherwise. Five temporal principal components of the BOLD signal time-courses extracted from the WM were regressed out of each voxel’s time-course; signal originating in the CSF was similarly regressed out. Six principal components corresponding to the six motion parameters estimated during offline motion correction were also regressed out, as well as their first time derivative. No low-pass filtering was applied.

### Modeling localizer data

For each localizer task, a general linear model estimated the effect size of each condition in each experimental run in each voxel. These effects were each modeled with a boxcar function (representing entire blocks) convolved with the canonical Hemodynamic Response Function (HRF). The model also included first-order temporal derivatives of these effects, as well as nuisance regressors representing entire experimental runs and offline-estimated motion parameters. The obtained beta weights were then used to compute the functional contrast of interest: Nonwords > Sentences for the MD localizer, and Sentences > Nonwords for the language localizer.

### Defining functional regions of interest (fROIs)

For each participant, functional ROIs were defined by combining two sources of information (following Fedorenko et al., 2010; Julian et al., 2012): (1) the participant’s activation map for the relevant localizer contrast, and (2) group-level spatial constraints (“masks”). The latter demarcated brain areas within which most or all individuals in prior studies showed activity for the localizer contrasts (**Figure 1**).

For the MD fROIs, we used masks derived from a group-level probabilistic representation of data from a previously validated MD-localizer task in a set of 197 participants. The task, described in detail in Fedorenko et al. (2011), contrasted hard and easy versions of a visuo-spatial working memory task (we did not use masks based on the Nonwords > Sentences contrast in order to maintain consistency with other current projects in our lab; prior work has established the similarity of the activation landscapes for these two contrasts, and the masks are sufficiently large such that slight differences in the activation landscapes, if they exist, wouldn’t affect our analyses; Fedorenko et al., 2013). These masks were constrained to be bilaterally symmetric by averaging individual Hard > Easy contrast maps across the two hemispheres prior to generating the group-level representation. The topography of these masks (available for download from http://web.mit.edu/evelina9/www/funcloc/funcloc_parcels.html) largely overlapped with anatomically based masks that were used in some prior studies (e.g. Fedorenko et al., 2013; Blank et al., 2014; Paunov et al., 2019). In particular, 10 masks were used in each hemisphere: in the posterior (PostPar), middle (MidPar), and anterior (AntPar) parietal cortex, precentral gyrus (PrecG), superior frontal gyrus (SFG), middle frontal gyrus (MFG) and its orbital part (MFGorb), opercular part of the inferior frontal gyrus (IFGop), the anterior cingulate cortex and pre-supplementary motor cortex (ACC/pSMA), and the insula (Insula).

For the language fROIs, we used masks derived from a group-level probabilistic representation for the Sentences > Nonwords contrast in a set of 220 participants. These masks (available for download from http://web.mit.edu/evelina9/www/funcloc/funcloc_parcels.html) were similar to the masks derived from 25 participants, as originally reported in Fedorenko et al. (2010), and covered extensive portions of the left lateral frontal and temporal cortex. In particular, six masks were used: three in the frontal lobe (in the inferior frontal gyrus (IFG), and its orbital part (IFGorb), and in the middle frontal gyrus (MFG)), and three in the temporal and parietal cortex (in the anterior temporal cortex (AntTemp), posterior temporal cortex (PostTemp), and in the angular gyrus (AngG)). We additionally defined the right hemisphere (RH) homologues of the language network regions. To do so, the left hemisphere (LH) masks were mirror-projected onto the RH to create six homologous masks.

The group-level masks, in the form of binary maps, were used to constrain the selection of subject-specific fROIs in each network. In particular, for each participant, 20 MD fROIs were created by intersecting each MD mask with that participant’s unthresholded *t*-map for the Nonwords > Sentences contrast; the 10% of voxels with the highest *t* values in the intersection image were then chosen as the fROI. In a parallel fashion, 12 language fROIs were created for each participant by intersecting each language mask with that participant’s unthresholded *t*-map for the Sentences > Nonwords contrast and selecting the 10% of voxels with the highest *t* values in the intersection image. A BOLD signal time-course for each story in the story listening task was then extracted from each voxel in each fROI of each participant.

We note that, for both the MD and language networks, only the group-based masks— which cover large swaths of cortex—were symmetric across hemispheres; the fROIs themselves were free to vary in their location between the two hemispheres, within the borders of these masks.

### Model of comprehension difficulty using self-paced reading times

To verify that self-paced reading times (SPRTs) reflect stimulus-related processing (following the logic in Hasson, Yang, Vallines, Heeger, & Rubin, 2008; Lerner, Honey, Silbert, & Hasson, 2011), we first computed inter-subject correlations among the time-series of per-word SPRTs for each story: each individual’s time-series was correlated with the average time-series of the rest of the participants. The average correlations varied between r=0.38 and r=0.59 across the stories and were all reliably above chance (all *p*s<10^−25^). As mentioned above, we used the default exclusion criteria used by (Futrell et al., 2018): we excluded data for a story if a participant answered less than 5 out of 6 questions wrong and outlier reading times of less than 100ms or more than 3,000ms were also excluded.

Mean reading times per word from the self-paced reading experiment (n=181) were temporally aligned with their corresponding word onsets in the auditory stimulus. Then, we obtained a per-TR time-series of SPRTs by averaging the reading times for the words that occurred within each TR (corresponding to 2 s) when the recorded stories were played in the fMRI experiment. Words that overlapped with two TRs were assigned to the TR with greater overlap. We then computed the average (across participants) per-TR SPRT, arriving at a final measure of comprehension difficulty at each TR.

### Model of comprehension difficulty using eye-tracking measures

We used four eye-tracking measures for participants in the eye-tracking study (n=40): (1) first pass regression (FPR), a variable indicating for each word whether or not a regressive eye-movement occurred from that word in the first pass; (2) regression path duration (RPD) or go-past time, the duration of the period between the onset of the first fixation on a word and the first fixation on anything to the right of it (RPD thus includes time spent on regressive fixations); (3) first pass reading time (FPRT) or gaze duration, the summed duration of all first-pass fixation durations on a word before any other word (left or right) is fixated; and (4) first fixation progressive (FFP), a variable indicating whether the first fixation on a word took place before any downstream words were viewed. To verify that eye-tracking measures reflect stimulus-related processing, we follow the same procedure as used for SPRTs, and compute inter-subject correlations among the time-series of per-word FPRs, RPDs, FPRTs and FFPs for each story. The average correlations varied between r=0.13 and r=0.17 across the stories for FPRs, between r=0.27 and r=0.42 across the stories for RPDs, between r=0.38 and r=0.53 across the stories for FPRTs, and between r=0.37 and r=0.53 across the stories for FFPs (all correlations higher than chance, all *p*s<10^−4^).

Mean eye-tracking measures per word were temporally aligned with their corresponding word onsets in the auditory stimulus. Then, we obtained a per-TR average measure of FPR, RPD, FPRT and FFP by averaging each of the four eye-tracking measures across the words that occurred within each 2s TR following the same procedure as used for SPRTs.

### Critical analysis using self-paced reading times and eye-tracking measures

Our analysis is summarized in **Figure 2**. As described above, for each TR *t*, we obtained SPRT, FPR, RPD, FPRT and FFP measures. We construct a design matrix for the experiment in which each row *t* corresponds to the concatenated five measures for a TR *t*. To account for the hemodynamic response, we model its effect as a fourth order finite impulse response (FIR) filter. We perform a simple linear regression: for each of the five variables, we estimate four weights that correspond to TRs *t*+1, *t*+2, *t*+3 and *t*+4 after onset and time *t*. Effectively, this corresponds to concatenating delayed versions of the design matrix so that each row *t* in the final design matrix contains the five measures for TRs *t*-4, *t*-3, *t*-2 and *t*-1. This is a common approach for accounting for the hemodynamic response (Huth et al., 2016; Wehbe et al., 2014), and the choice of an 8s window is typically used to capture the effect of stimulus features on the fMRI response. This encoding model analysis (Huth et al., 2016; Wehbe et al., 2014) differs from the typical GLM analysis in two ways. First, instead of assuming a fixed hemodynamic response function that is constant across the brain, this approach allows for variability in the hemodynamic response by implicitly estimating it at each voxel. And second, instead of running a significance test on the regression weights, we run a more stringent test: we assess the generalization and stability of the learned weights by using them to predict held-out fMRI data unseen in training.

In particular, for each participant, we estimated generalization via a cross-validation scheme in which we iteratively held out one story and learned the regression weights on the remaining stories. We then predicted BOLD activity for the held-out story using the (delayed) design matrix for that story and the learned regression weights. This procedure resulted in a predicted time series of BOLD activity in each voxel in each fROI of each participant during the held-out story. We then measured how closely these predictions correspond to the real data via Pearson’s correlation. This correlation was computed between the average (across voxels) fROI activity predicted by the model and the corresponding average fROI activity in the real data to obtain summary statistics for the fROIs. Finally, we averaged the correlation values across all cross-validation folds to obtain a single correlation value per fROI per participant (i.e., 42 mean correlation values for each fROI, one for each participant).

To better characterize the findings at the level of the networks of interest, the above analysis was repeated, but this time, predicted and actual BOLD time series were grouped into four sets: Left Hemisphere (LH) Language fROIs, Right Hemisphere (RH) Language fROIs, LH MD fROIs, and RH MD fROIs.

It is worth mentioning that the direction of prediction we used here (predicting brain activity from comprehension difficulty measures instead of the other way around) was in keeping with the encoding model approach (Huth et al., 2016; Naselaris, Kay, Nishimoto, & Gallant, 2011; Wehbe et al., 2014) and does not imply that brain activity is caused by comprehension difficulty measures. Typically, when using encoding models with stimulus features to predict brain activity, one can state causal statements unambiguously (Weichwald et al., 2015). However, here we use the encoding approach as a way to only measure correlation between two effects (fMRI activity and reading times) of the same cause (comprehension difficulty).

### Noise ceiling correction

To help with interpreting prediction performance, we provide measures of prediction performance that are corrected by the estimated noise ceiling for each fROI and fROI group. The noise ceiling is an approximation of the maximum possible performance. fMRI stimuli engage brain regions to a different extent, and regions have different physiological characteristics, both of which affect the signal-to-noise ratio. We estimate the noise ceiling for each fROI (and fROI group) across participants by adapting the method proposed by Hsu et al. (2004) to be used for multiple participants. To evaluate the noise ceiling, Hsu et al. (2004) consider different repeats of the same stimulus that is seen by multiple participants. We treat the average fROI activity for the subjects listening to the same story as different repeats of the same story. For each story and each fROI, we evaluate the noise ceiling by first computing the average time-course of this fROI for each of the k subjects that have listened to this story. We then compute the correlation of each of the 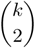 pairs of time series. We then average all these pairwise correlations, and further average these estimates for all the stories. We end up with a measure of noise ceiling for each fROI. We repeat this approach for fROI groups. Following previous work (A. Hsu et al., 2004; Lescroart & Gallant, 2019; Lescroart, Stansbury, & Gallant, 2015), we normalize the average prediction performance by the square-root of the noise-ceiling, yielding normalized correlation values.

### Computing confidence intervals

The participant-specific (unnormalized and normalized) correlation values were averaged across participants, and empirical confidence intervals were estimated for the mean prediction in each fROI, by running a bootstrap test. Here are the details: the following was repeated 50,000 times: a set of 42 participants was sampled with replacement from the original 42 participants, and correlation values of the sampled 42 participants were averaged for each fROI. 90, 95, 98, and 99% confidence intervals were constructed from the 50,000 samples. Finally, for each fROI, the smallest confidence interval containing 0 was transformed into a *p*-value and the Holm-Bonferroni method was applied to correct for multiple hypothesis testing (Holm, 1979). In this context where we have a relatively low number of hypotheses, controlling for the family-wise error rate (e.g. by using Holm-Bonferroni) is more appropriate than controlling the false discovery rate. Since normalized correlations are a rescaling of the unnormalized correlations, we apply a single test for the unnormalized correlations (with the results of the normalized correlations being the same).**Comparing the two networks.** To evaluate whether the average correlation across participants is higher in the Language network than in the MD network we again ran a bootstrap test. First, we computed for each subject the difference between the average correlation in the LH Language and MD fROI group. This resulted in 42 values. We then repeated the following 50,000 times: a set of 42 participants was sampled with replacement from the original 42 participants, and the LH difference for the 42 sampled participants is averaged. This set of 50,000 samples yielded an empirical distribution from which we computed a *p*-value. We repeated this test to obtain a *p*-value for the difference between the RH Language and MD fROI group. We included these two *p*-values in the multiple testing correction mentioned in the previous section.

## Results

The effect size of the unnormalized correlations between online comprehension difficulty and BOLD activity in the networks of interest is small, in part because of the low signal-to-noise ratio of fMRI. For this reason, we interpreted the size of this effect by taking into account a metric measure of signal reliability based on inter-subject correlation of the fMRI signals, effectively performing a noise ceiling correction (see Methods) (Blank & Fedorenko, 2017; A. Hsu et al., 2004; Lescroart & Gallant, 2019; Lescroart et al., 2015). This choice of a ceiling metric leads to conservative normalized correlations: Blank and Fedorenko (2017) show that within-subject correlations are lower than cross-subject correlations on this task, and would consequently lead to larger normalized correlations.

Self-paced reading times and eye-tracking measures predicted BOLD activity in the language network, based on either unnormalized or normalized correlations. Average (normalized) prediction performance is significantly greater than chance (false discovery rate controlled at 0.01) in both the LH and RH language network (**Figure 3a**), including in most individual fROIs (the bilateral IFGorb, IFG, MFG, AntTemp, and PostTemp fROIs, and the left AngG fROI; **Figure 3b**). In contrast, comprehension difficulty measures did not significantly predict activity in the MD network, either when averaging across fROIs within the LH or RH, or in any individual MD fROI. (It is worth noting that we chose to run the significance tests on the unnormalized correlations but running it on the normalized correlations, shown in **Figure 3c-d**, would lead to a similar result since the intervals are normalized by a constant, as can be judged by the similarity of the confidence intervals.)

**Table 2:**
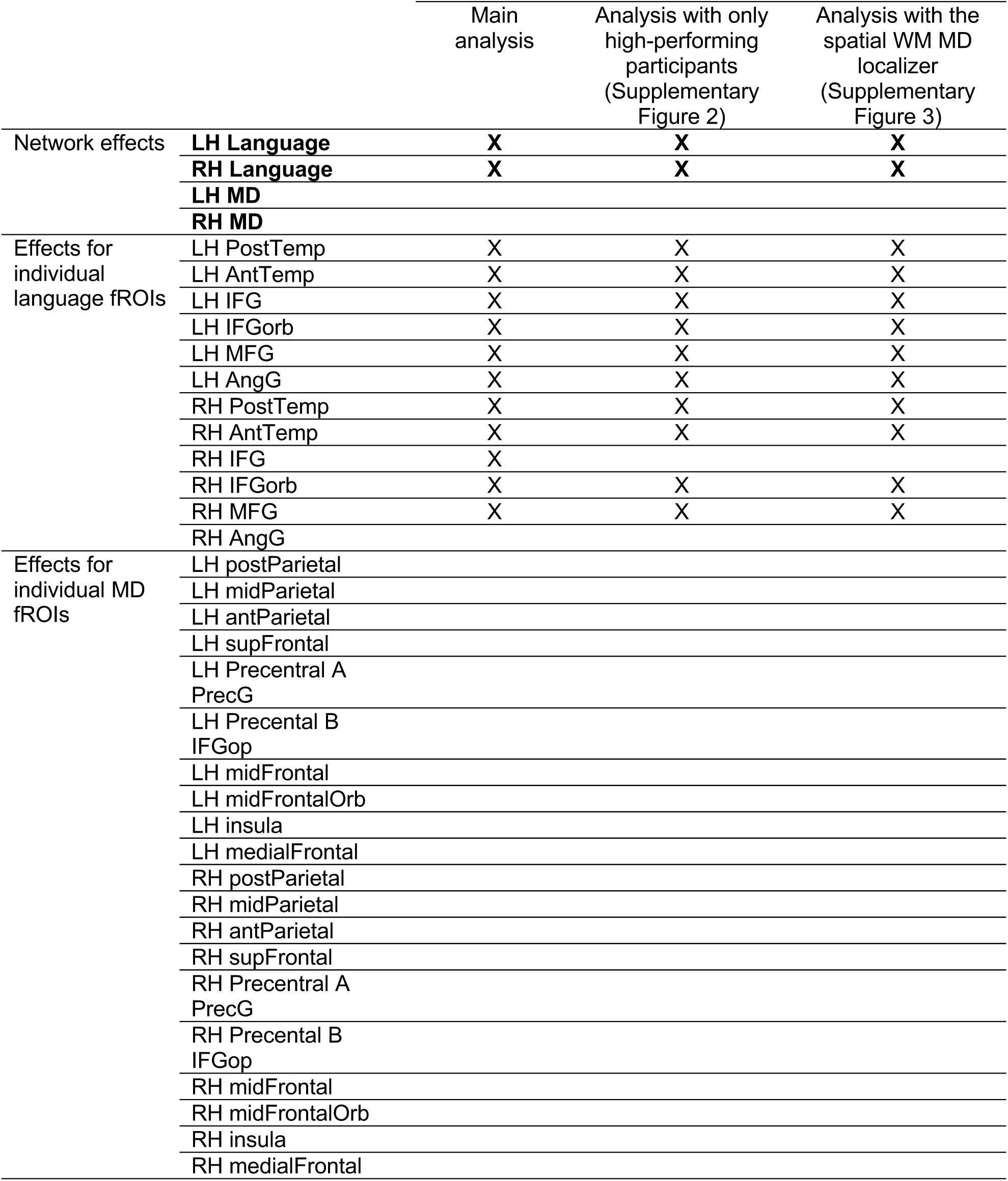
Summary of results across fROI groups (bolded) and fROIs, for the main analysis and the two additional supplementary analyses. X marks a significantly higher than 0 correlation. The three sets of results are extremely similar, except for a small variation in one fROI in the last set.

To directly compare between the language and the MD network, we estimated a *p*-value for a two-sample test by first computing the difference between the prediction performance in the language and MD networks for each subject and then using a bootstrap procedure. We find that the average unnormalized correlation for the language network in each hemisphere is significantly larger than the unnormalized correlation for the MD network (*p*= 2×10^−5^ for LH and *p*=2×10^−5^ for RH). False discovery rate was controlled at 0.01.

As described in the Methods, we additionally performed two other versions of this analysis: one, where we only included participants with near-perfect accuracies on the comprehension questions for the story listening task, and one, where we used a visuo-spatial working memory task to localizer the MD regions (which also corresponded to a smaller set of participants). The results of both of these additional analyses mirrored the results of the main analysis (with a small exception for the correlation in the RH IFG not being significantly higher than zero), where the original study included all participants and used the Nonwords > Sentences contrast to localize the MD regions (see **Table 3** and **Supplementary Figures 2** and **3**).

## Discussion

In this study, we tested whether language comprehension—in addition to language-specific resources—recruits domain-general executive mechanisms. To this end, we examined the relationship between behavioral and neural measures of incremental comprehension difficulty during naturalistic language processing: behavioral comprehension difficulty was estimated with two commonly used approaches (self-paced reading times and eye-tracking fixation durations); and neural recruitment was quantified from fMRI BOLD activity in domain-general and language-specific functional networks that have been previously implicated in language comprehension. We found that, whereas neural activity in the fronto-temporal, language-selective network (Braga, DiNicola, & Buckner, 2019; Fedorenko et al., 2011; Fedorenko & Thompson-Schill, 2014) was predicted by behavioral measures of incremental comprehension difficulty, activity in the domain-general MD network (Duncan, 2010) was not predicted by these measures. Furthermore, this difference between networks was reliable: the mean prediction performance was significantly higher in the LH or RH language network than in the LH and RH MD network, respectively. The lack of a reliable correlation between behavioral comprehension difficulty and neural activity in the MD network conflicts with previous studies reporting MD activity during some language tasks and its sensitivity to linguistic manipulations of processing difficulty (January, Trueswell, & Thompson-schill, 2009; Kuperberg et al., 2003; McMillan et al., 2013; Nieuwland et al., 2012; Novais-Santos et al., 2007; Peelle et al., 2010; Rodd et al., 2005).

Our investigation complements a prior study, Henderson et al. (2015), which related behavioral and neural measures of language comprehension by co-registering eye-tracking and fMRI data during naturalistic reading, using fixation durations to predict neural activity. Because behavioral and neural measures were obtained from the same participants, an experimental control (a pseudo-reading task) was used in order to isolate linguistic components of the signal. Activity in parts of the left middle and superior temporal gyri was found to correlate more strongly with fixation durations during the reading of natural text compared to a perceptually matched meaningless control materials, suggesting that longer fixations were at least partially driven by effort related to linguistic processing in brain areas commonly associated with language comprehension, i.e., putative parts of the “core language network”. Similar to our study, Henderson et al. (2015) did not observe a correlation between their comprehension difficulty measure and activity in the fronto-parietal MD regions. However, they adopt a traditional group-based analytic approach, which relies on voxel-wise correspondence across individuals and does not take into account the well-known inter-individual variation in the precise locations of functional areas in the association cortex (e.g. Fischl et al., 2008; Frost & Goebel, 2012; Tahmasebi et al., 2012; Vázquez-Rodríguez et al., 2019). Because of the resulting low sensitivity of such analyses (e.g. Nieto-Castañón & Fedorenko, 2012), Henderson et al. (2015) may have missed the effects within the MD network. In the current study, to maximize the probability of detecting a relationship between behavioral measures and MD activity, we functionally localized MD areas (as well as core language areas) in each individual participant (Fedorenko et al., 2010). This strategy reduces the risk of obtaining a false negative for the MD network.

Our use of separate participant pools in the behavioral studies vs. the fMRI study further eliminates many non-linguistic confounds (such as attention and motor control processes related to eye-movements or button presses) that are difficult to avoid when the behavioral and neural measures come from the same individuals (e.g., Henderson et al., 2015). The observed relationship between reading latencies and brain activity is thus most plausibly due to *linguistic properties* of the story stimuli. These findings support the hypotheses that (1) behavioral responses to language stimuli reflect computational load in language comprehension mechanisms (Just & Carpenter, 1980); (2) language comprehension difficulty generalizes across participants, task demands, and modality of presentation (Hasson et al., 2008); and (3) processing mechanisms that give rise to measurable reading delays reside in a language-selective cortical network, rather than in a domain-general executive control network. In this way, our results reinforce those of Henderson et al. (2015): we replicate their finding of a relationship between reading latencies and neural activity in the left-hemisphere middle and superior temporal lobe, and extend it to other parts of the language network, including the language-responsive areas in the left inferior frontal cortex and, to a lesser extent, the left angular gyrus, as well as the right-hemisphere homologs of the left-hemisphere language regions. These more widespread effects across the language network are plausibly due to increased sensitivity resulting from participant-specific functional localization.

At present, much behavioral language research is disconnected from cognitive neuroscience efforts to understand the architecture of language comprehension, despite (1) the fact that these two enterprises share the same goal—to understand the computations that support language comprehension, and (2) the fact that a link between behavioral measures of language comprehension, or the mental states they correspond to, and neural correlates of language comprehension is a fundamental assumption of psycholinguistics (e.g. Just & Carpenter, 1980). Indeed, except for Henderson et al. (2015) and the current paper, cognitive neuroscientists have not typically used direct and continuous behavioral measures to model brain activity during language comprehension (see e.g., **Supplementary Table 1** for fMRI studies that have used naturalistic linguistic materials and which have typically used linguistic features as predictors of neural activity, often without first establishing a link between those features and *behavioral* measures). The current paper connects the psycholinguistic and cognitive neuroscience literatures, and in so doing contributes to both fields. For psycholinguistics, our results validate widely used behavioral measures as indeed revealing the underlying activity of language comprehension mechanisms. For cognitive neuroscience, our results indicate that, even using a broad (all-encompassing) and theory-neutral estimate of comprehension difficulty, language processing recruits primarily cortical circuits that specialize for this purpose, and that domain-general executive mechanisms are generally not recruited during naturalistic sentence comprehension.

This work thus sheds new light on the role of the domain-general MD network in language comprehension, and on the division of labor between these domain-general mechanisms and the language-selective ones. In particular, regions of the MD network have been shown to be sensitive to linguistic difficulty across diverse manipulations (see Fedorenko (2014), for a review). However, almost all prior evidence has come from traditional, task-based experimental paradigms that present participants with linguistic manipulations that do not commonly occur in real-life comprehension scenarios (like ambiguous words that are not disambiguated by the context, or non-local dependencies; e.g., January, Trueswell, & Thompson-Schill, 2009; Novais-Santos et al., 2007; Peelle, Troiani, Wingfield, & Grossman, 2010; Rodd et al., 2005) and ask them to solve “artificial” tasks, such as making similarity judgments or deciding whether a sentence matches a picture. Although the stories used in the current study were modified to include challenging linguistic phenomena in order to increase variability in processing demands and increase the chances of engaging executive resources, the only “task” required of participants was naturalistic comprehension of the narratives. The fact that we do not find a relationship between comprehension difficulty and the MD network’s activity in our study suggests that the MD network’s contribution to language comprehension may be restricted to artificial scenarios, where language is effectively turned into problem solving (Diachek et al., in press; P. Wright, Randall, Marslen-Wilson, & Tyler, 2011). In line with this conjecture, Blank and Fedorenko (2017) showed that MD regions do not strongly track language stimuli during comprehension, and Shain and colleagues (2019) showed that activity in the MD regions during comprehension does not correlate with the psycholinguistic construct of “surprisal”, the moment-by-moment unpredictability of linguistic input (see also Blanco-Elorrieta & Pylkkänen, 2017, for evidence of less MD engagement during a more naturalistic production paradigm). Whereas the MD network may play *some* role during language processing (perhaps modulating overall alertness or attention) our results as well as others mentioned above suggest that this system is not directly involved in linguistic computations.

Our results also suggest that similarities between language processing and other kinds of processing (e.g., theoretical constructs in mathematics or music resembling those in natural language syntax) do not entail shared neural circuitry (see also Fedorenko & Blank, 2020, for a recent discussion). In particular, the fact that multiple domains require hierarchical combinatoric processing of symbols does not mean that the same circuits are engaged across these domains.

Rather, constructing hierarchical sequences, predictive coding, working memory storage and retrieval of information, and other processes that may be necessary in multiple domains of cognition appear to be implemented within domain-specialized systems, including the language processing areas.

In conclusion, we found that whereas neural activity in the fronto-temporal language network is predicted by behavioral signatures of incremental comprehension difficulty, activity in in the domain-general fronto-parietal multiple demand network is not.

## Author contributions

LW, IB, EG, and EF designed research. All authors performed research: LW, IB, and EF collected, preprocessed, and analyzed fMRI data; RF, IB, and EG collected, preprocessed, and analyzed the self-paced reading data; RL, TM, and NS collected, preprocessed, and analyzed the eye-tracking data. All authors interpreted the data. LW and EF wrote the manuscript. IB, CS, RF, RL, TM, and EG provided comments.

## Acknowledgments

LW was supported by startup funds at Carnegie Mellon University. RF was supported by NSF DDRI grant number 1551543. TM was supported by a Feodor Lynen Research Fellowship awarded to him by the Alexander von Humboldt Foundation and by NIH grant HD065829 awarded to RL and Keith Rayner. RL was also supported by an Alfred P. Sloan Research Fellowship and a Paul and Lilah Newton Brain Science Award. EF was supported by NIH grants HD057522, DC016607, and DC016950, a grant from the Simons Foundation to the Simons Center for the Social Brain at MIT, and by the Department of Brain and Cognitive Sciences and McGovern Institute for Brain Research at MIT. We acknowledge the Athinoula A. Martinos Imaging Center at the McGovern Institute for Brain Research, MIT. For technical support during scanning, the authors thank Steve Shannon, Christina Triantafyllou, and Atsushi Takahashi. We thank Anastasia Vishnevetsky and Steve Piantadosi for help in constructing the story materials, Nancy Kanwisher for help in recording the story materials, Kris Fedorenko for editing the recordings, Zach Mineroff for help in collecting the fMRI data and preparing the data for analyses, Hal Tily for help in collecting the self-paced reading data, Kyle Mahowald for help in preprocessing the reading data, Wade Shen and Jeanne Gallée for help in aligning the story transcripts to the auditory files.

---

**Supplementary Table 1:**
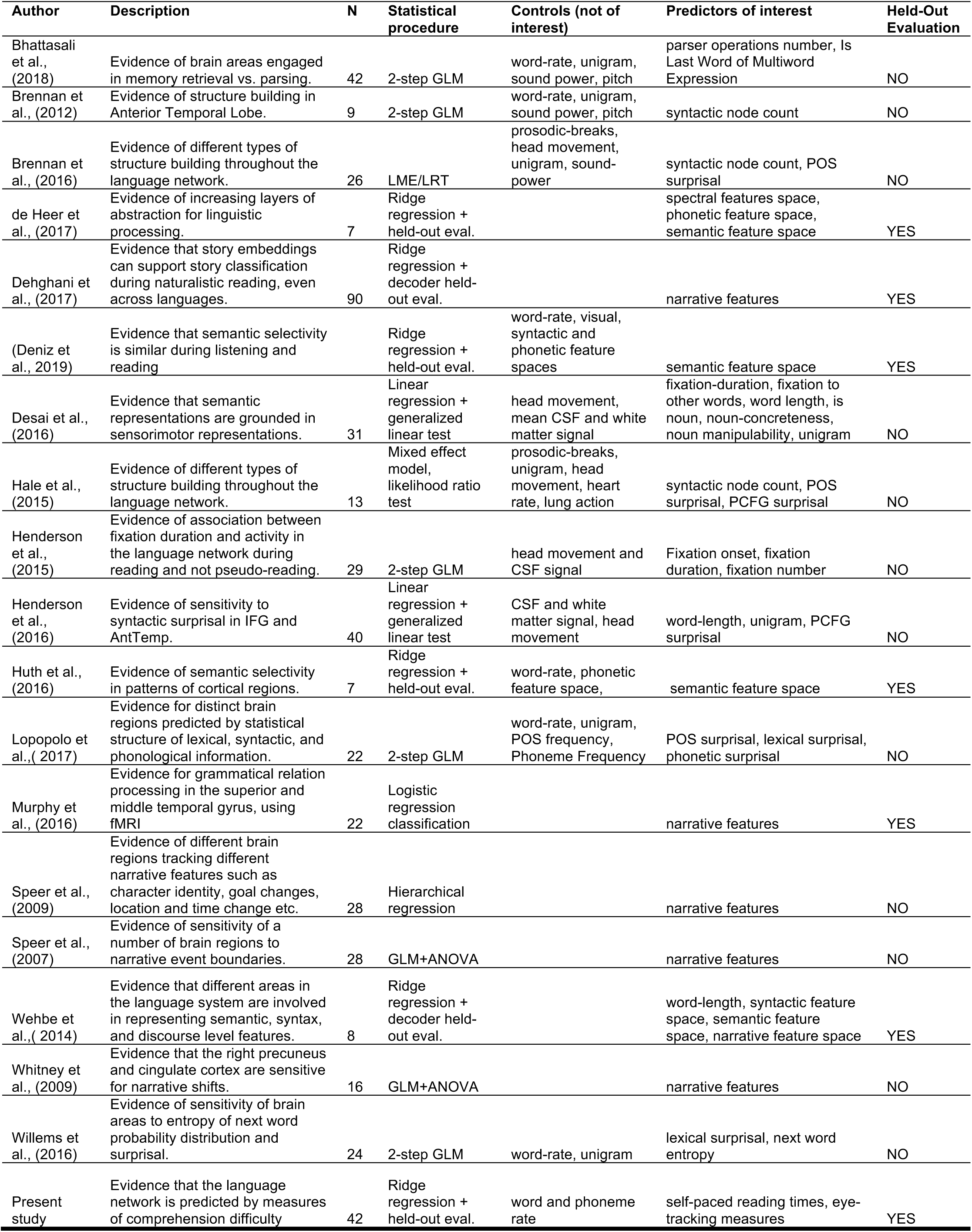
Studies that used naturalistic linguistic materials with the goal

**Supplementary Figure 1.**
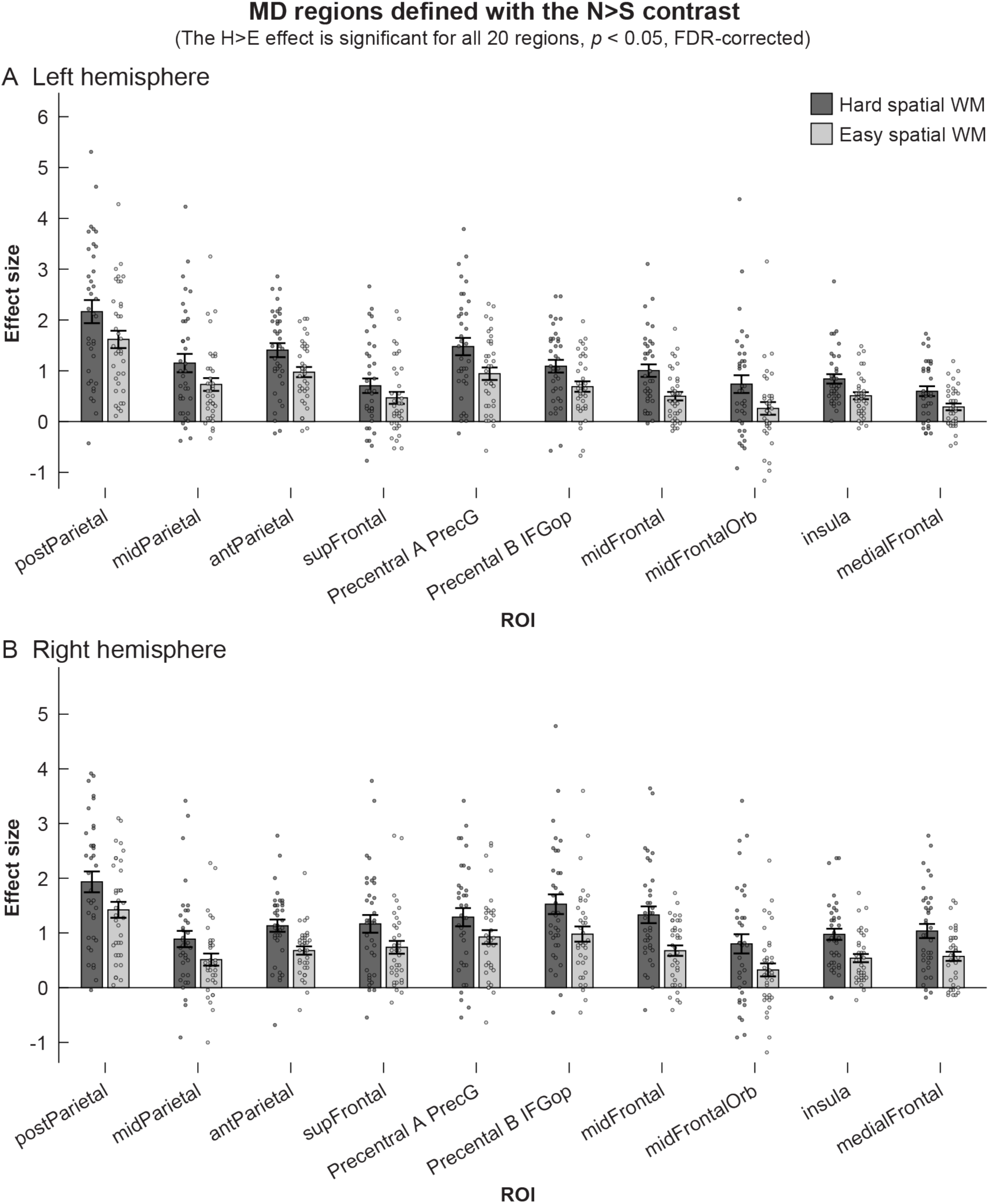
Response of MD regions defined with the Nonwords > Sentences contrast to the Hard and Easy conditions of the visuo-spatial working memory MD localizer.

**Supplementary Figure 2.**
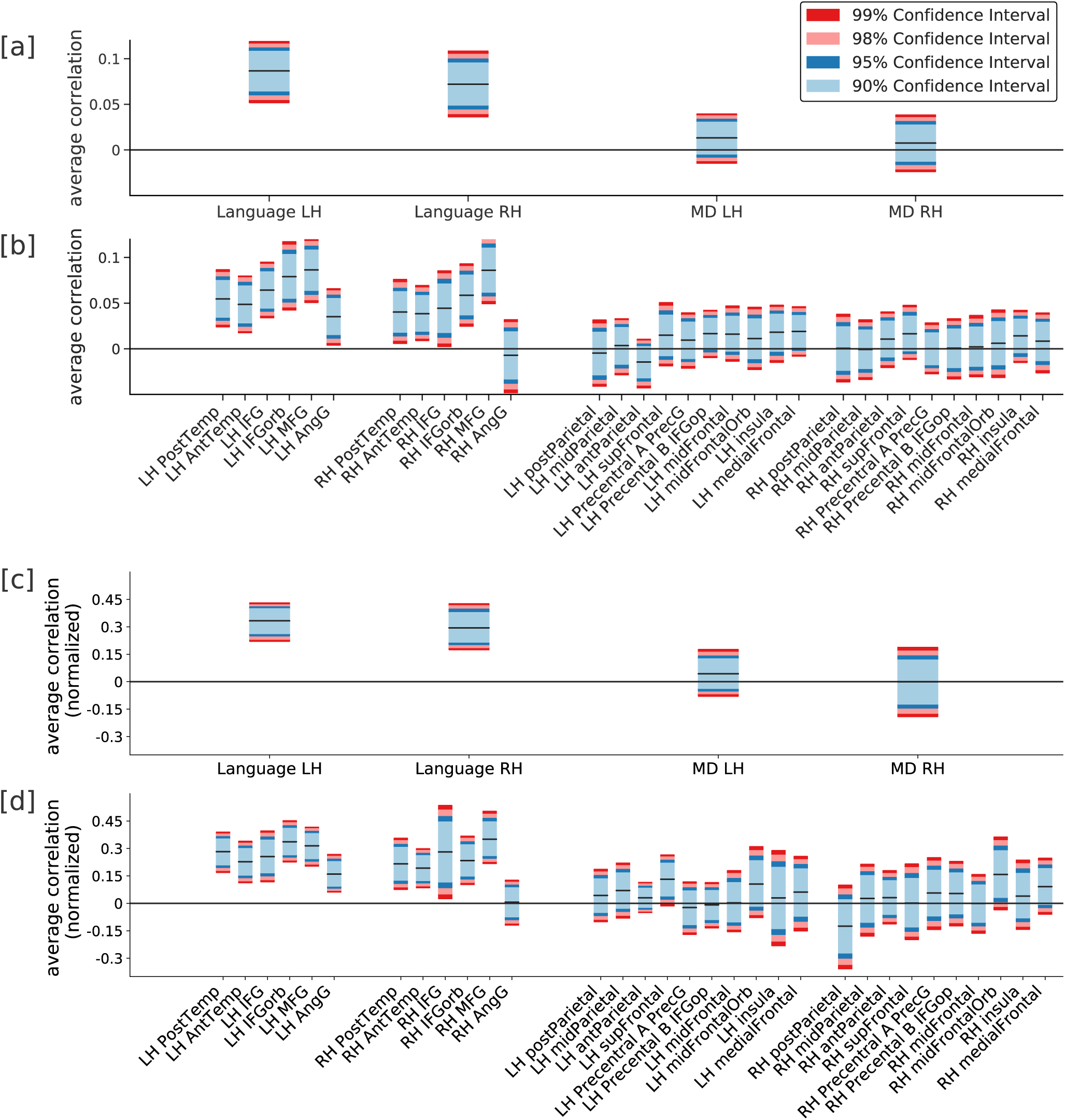
Average (unnormalized and normalized) correlation between activity predicted as a function of comprehension difficulty (estimated using self-paced reading times and eye-tracking measures) and real held-out activity, normalized by the estimated reliability of the signal for each fROI group ([a] unnormalized and [c] normalized) and each fROI ([b] unnormalized and [d] normalized). Performance was averaged across the 42 participants and bootstrap confidence intervals were constructed. Reading times predict the activity in left and right language fROIs, but not in MD fROIs. **Here the MD fROIs were defined using the spatial MD localizer and the results are shown only for the subset of 35 subjects that underwent this task.**

**Supplementary fig. 3.**
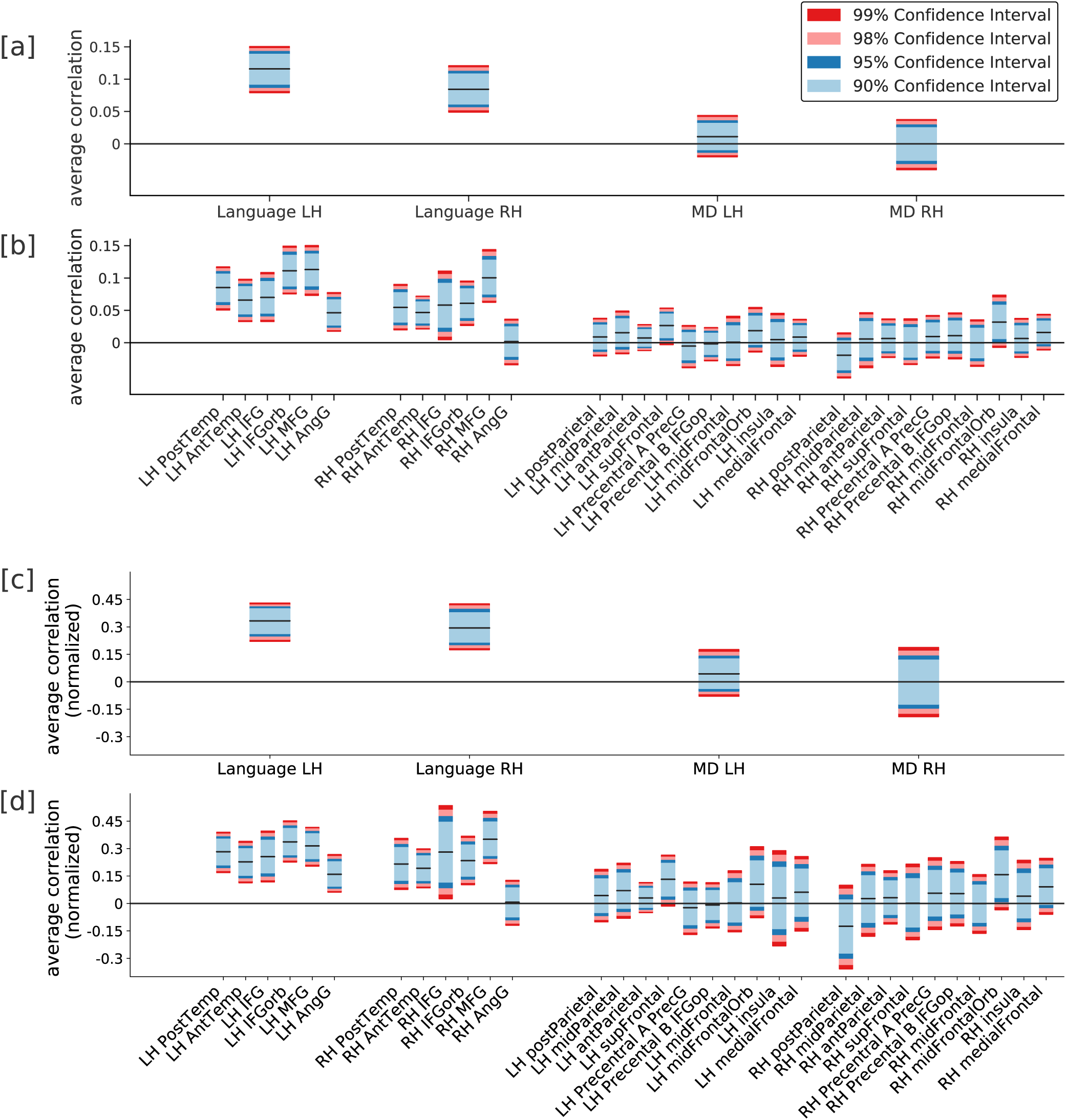
Average (unnormalized and normalized) correlation between activity predicted as a function of comprehension difficulty (estimated using self-paced reading times and eye-tracking measures) and real held-out activity, normalized by the estimated reliability of the signal for each fROI group ([a] unnormalized and [c] normalized) and each fROI ([b] unnormalized and [d] normalized). **The analysis is restricted here to the 24 subjects with the best performance.** Performance was averaged across these 24 participants and bootstrap confidence intervals were constructed. Reading times predict the activity in left and right language fROIs, but not in MD fROIs.

## Footnotes

1 The Cohen correction is designed to correct for poor eye-tracker calibration. However, poor calibration is reflected in fixation offsets in the same direction, and variable correction vectors therefore indicate that the Cohen correction failed.

## References

Aaronson, D., & Scarborough, H. S. (1977). Performance theories for sentence coding: Some quantitative models. Journal of Verbal Learning and Verbal Behavior, 16(3), 277–303. https://doi.org/10.1016/S0022-5371(77)80052-2

Abney, S. P., & Johnson, M. (1991). Memory requirements and local ambiguities of parsing strategies. Journal of Psycholinguistic Research, 20(3), 233–250.

Anderson, M. L. (2010). Neural reuse: A fundamental organizational principle of the brain. Behavioral and Brain Sciences. https://doi.org/10.1017/S0140525×10000853

Ayyash, D.*, Malik Moraleda, S. M.., Galleé, J. M. Z., Jouravlev, O., & Fedorenko, E. (n.d.). The universal language network: A cross-linguistic investigation spanning 41 languages and 10 language families. In Preparation.

Bautista, A., & Wilson, S. M. (2016). Neural responses to grammatically and lexically degraded speech. Language, Cognition and Neuroscience, 31(4), 567–574.

Bhattasali, S., Hale, J., Pallier, C., Brennan, J. R., Luh, W.-M., & Spreng, R. N. (2018). Differentiating Phrase Structure Parsing and Memory Retrieval in the Brain. Proceedings of the Society for Computation in Linguistics (SCiL) 2018, Salt Lake City, Utah, January 4-7, 2018, (2012), 74–80. https://doi.org/10.7275/R5FF3QJ2

Bilenko, N. Y., Grindrod, C. M., Myers, E. B., & Blumstein, S. E. (2008). Neural correlates of semantic competition during processing of ambiguous words. Journal of Cognitive Neuroscience, 21(5), 960–975.

Binder, J. R. (1997). Neuroanatomy of language processing studied with functional MRI. Clinical Neuroscience.

Blanco-Elorrieta, E., & Pylkkänen, L. (2017). Bilingual Language Switching in the Laboratory versus in the Wild: The Spatiotemporal Dynamics of Adaptive Language Control. Journal of Neuroscience, 37(37), 9022–9036. https://doi.org/10.1523/JNEUROSCI.0553-17.2017

Blank, I., Balewski, Z., Mahowald, K., & Fedorenko, E. (2016). Syntactic processing is distributed across the language system. Neuroimage, 127, 307–323.

Blank, I., & Fedorenko, E. (2017). Domain-general brain regions do not track linguistic input as closely as language-selective regions. The Journal of Neuroscience, 3642–16. https://doi.org/10.1523/JNEUROSCI.3642-16.2017

Blank, I., Kanwisher, N., & Fedorenko, E. (2014). A functional dissociation between language and multiple-demand systems revealed in patterns of BOLD signal fluctuations. Journal of Neurophysiology, 112(5), 1105–1118. https://doi.org/10.1152/jn.00884.2013

Blumstein, S. E., & Amso, D. (2013). Dynamic Functional Organization of Language: Insights From Functional Neuroimaging. Perspectives on Psychological Science. https://doi.org/10.1177/1745691612469021

Braga, R. M., DiNicola, L. M., & Buckner, R. L. (2019). Situating the Left-Lateralized Language Network in the Broader Organization of Multiple Specialized Large-Scale Distributed Networks. BioRxiv. https://doi.org/10.1101/2019.12.11.873174

Braze, D., Mencl, W. E., Tabor, W., Pugh, K. R., Constable, R. T., Fulbright, R. K., … Shankweiler, D. P. (2011). Unification of sentence processing via ear and eye: An fMRI study. Cortex, 47(4), 416–431.

Brennan, J. R., Stabler, E. P., Van Wagenen, S. E., Luh, W. M., & Hale, J. T. (2016). Abstract linguistic structure correlates with temporal activity during naturalistic comprehension. Brain and Language, 157–158, 81–94. https://doi.org/10.1016/j.bandl.2016.04.008

Brennan, J, Nir, Y., Hasson, U., Malach, R., Heeger, D. J., Pylkkänen, L., … Structures, T. (2010). Syntactic structure building in the anterior temporal lobe during natural story listening. Brain and Language, 6(2), 247–253. https://doi.org/10.1111/j.1743-6109.2008.01122.x.Endothelial

Brennan, Jonathan, Nir, Y., Hasson, U., Malach, R., Heeger, D. J., & Pylkkänen, L. (2012). Syntactic structure building in the anterior temporal lobe during natural story listening. Brain and Language, 120(2), 163–173. https://doi.org/10.1016/j.bandl.2010.04.002

Brennan, Jonathan, & Pylkkänen, L. (2012). The time-course and spatial distribution of brain activity associated with sentence processing. NeuroImage, 1–10. https://doi.org/10.1016/j.neuroimage.2012.01.030

Broca, P. P. (1861). Remarks on the seat of the faculty of articulated language, following an observation of aphemia (loss of speech). Bulletin de La Société Anatomique.

Campbell, K. L., & Tyler, L. K. (2018). Language-related domain-specific and domain-general systems in the human brain. Current Opinion in Behavioral Sciences. https://doi.org/10.1016/j.cobeha.2018.04.008

Clifton, C., & Frazier, L. (1989). Comprehending Sentences with Long-Distance Dependencies BT - Linguistic Structure in Language Processing. In G. N. Carlson & M. K. Tanenhaus (Eds.) (pp. 273–317). Dordrecht: Springer Netherlands. https://doi.org/10.1007/978-94-009-2729-2_8

Cohen, A. L. (2013). Software for the automatic correction of recorded eye fixation locations in reading experiments. Behavior Research Methods, 45(3), 679–683.

Cohen, J., Cohen, P., West, S. G., & Aiken, L. S. (2013). Applied multiple regression/correlation analysis for the behavioral sciences. Routledge.

D’Esposito, M., & Postle, B. R. (2015). The Cognitive Neuroscience of Working Memory. Annual Review of Psychology. https://doi.org/10.1146/annurev-psych-010814-015031

de Heer, W. A., Huth, A. G., Griffiths, T. L., Gallant, J. L., & Theunissen, F. E. (2017). The Hierarchical Cortical Organization of Human Speech Processing. The Journal of Neuroscience, 37(27), 6539–6557. https://doi.org/10.1523/JNEUROSCI.3267-16.2017

Dehghani, M., Boghrati, R., Man, K., Hoover, J., Gimbel, S. I., Vaswani, A., … Kaplan, J. T. (2017). Decoding the neural representation of story meanings across languages. Human Brain Mapping, 38(12), 6096–6106. https://doi.org/10.1002/hbm.23814

Demberg, V., & Keller, F. (2008). Data from eye-tracking corpora as evidence for theories of syntactic processing complexity. Cognition, 109(2), 193–210.

Deniz, F., Nunez-Elizalde, A. O., Huth, A. G., & Gallant, J. L. (2019). The Representation of Semantic Information Across Human Cerebral Cortex During Listening Versus Reading Is Invariant to Stimulus Modality. The Journal of Neuroscience: The Official Journal of the Society for Neuroscience. https://doi.org/10.1523/JNEUROSCI.0675-19.2019

Desai, R. H., Choi, W., Lai, V. T., & Henderson, J. M. (2016). Toward Semantics in the Wild: Activation to Manipulable Nouns in Naturalistic Reading. Journal of Neuroscience, 36(14), 4050–4055. https://doi.org/10.1523/JNEUROSCI.1480-15.2016

Diachek, E., Blank, I., Siegelman, M., Affourtit, J., & Fedorenko, E. (in press). The domain-general multiple demand (MD) network does not support core aspects of language comprehension: a large-scale fMRI investigation. https://doi.org/10.1101/744094

Dronkers, N. F., Wilkins, D. P., Van Valin, R. D., Redfern, B. B., & Jaeger, J. J. (2004). Lesion analysis of the brain areas involved in language comprehension. Cognition, 92(1–2), 145–177. https://doi.org/10.1016/j.cognition.2003.11.002

Duncan, J. (2010). The multiple-demand (MD) system of the primate brain: mental programs for intelligent behaviour. Trends in Cognitive Sciences, 14(4), 172–179. https://doi.org/10.1016/j.tics.2010.01.004

Eklund, A., Andersson, M., Josephson, C., Johannesson, M., & Knutsson, H. (2012). Does parametric fMRI analysis with SPM yield valid results?-An empirical study of 1484 rest datasets. NeuroImage. https://doi.org/10.1016/j.neuroimage.2012.03.093

Fadiga, L., Craighero, L., & D’Ausilio, A. (2009). Broca’s area in language, action, and music. In Annals of the New York Academy of Sciences. https://doi.org/10.1111/j.1749-6632.2009.04582.x

Fedorenko, E., Behr, M. K., & Kanwisher, N. (2011). Functional specificity for high-level linguistic processing in the human brain. Proceedings of the National Academy of Sciences, 108(39), 16428–16433. https://doi.org/10.1073/pnas.1112937108

Fedorenko, E, Hsieh, P.-J., Nieto-Castanon, A., Whitfield-Gabrieli, S., & Kanwisher, N. (2010). New method for f{MRI} investigations of language: Defining {ROI}s functionally in individual subjects. Journal of Neurophysiology, 104(2), 1177–1194.

Fedorenko, E, Nieto-Castanon, A., & Kanwisher, N. (2012). Lexical and syntactic representations in the brain: An f{MRI} investigation with multi-voxel pattern analyses. Neuropsychologia, 50(4), 499–513.

Fedorenko, E. (in press). The brain network that supports high-level language processing. In Gazzaniga, Ivry Mangun (Ed.), Cognitive Neuroscience: The Biology of the Mind (5th Edition).

Fedorenko, E. (2014). The role of domain-general cognitive control in language comprehension. Frontiers in Psychology, 5.

Fedorenko, E, & Blank, I. A. (2020). Broca’s Area Is Not a Natural Kind. Trends in Cognitive Sciences. https://doi.org/10.1016/j.tics.2020.01.001

Fedorenko, E, Blank, I., Siegelman, M., & Mineroff, Z. (2020). Lack of selectivity for syntax relative to word meanings throughout the language network. BioRxiv, 477851.

Fedorenko, E, Duncan, J., & Kanwisher, N. (2013). Broad domain generality in focal regions of frontal and parietal cortex. Proceedings of the National Academy of Sciences, 110(41), 16616–16621. https://doi.org/10.1073/pnas.1315235110

Fedorenko, E, Gibson, E., & Rohde, D. (2006). The nature of working memory capacity in sentence comprehension: Evidence against domain-specific working memory resources. Journal of Memory and Language. https://doi.org/10.1016/j.jml.2005.12.006

Fedorenko, E, Gibson, E., & Rohde, D. (2007). The nature of working memory in linguistic, arithmetic and spatial integration processes. Journal of Memory and Language. https://doi.org/10.1016/j.jml.2006.06.007

Fedorenko, E, Hsieh, P.-J., Nieto-Castanon, A., Whitfield-Gabrieli, S., & Kanwisher, N. (2010). New Method for fMRI Investigations of Language: Defining ROIs Functionally in Individual Subjects. Journal of Neurophysiology, 104(2), 1177–1194. https://doi.org/10.1152/jn.00032.2010

Fedorenko, E, Mineroff, Z., Siegelman, M., & Blank, I. (2018). Word meanings and sentence structure recruit the same set of fronto-temporal regions during comprehension. BioRxiv, 477851.

Fedorenko, E, Scott, T. L., Brunner, P., Coon, W. G., Pritchett, B., Schalk, G., & Kanwisher, N. (2016). Neural correlate of the construction of sentence meaning. Proceedings of the National Academy of Sciences, 113(41), E6256--E6262. https://doi.org/10.1073/pnas.1612132113

Fedorenko, E, & Thompson-Schill, S. L. (2014). Reworking the language network. Trends in Cognitive Sciences, 18(3), 120–127. https://doi.org/10.1016/j.tics.2013.12.006

Fedorenko, E, & Varley, R. (2016). Language and thought are not the same thing: Evidence from neuroimaging and neurological patients. Annals of the New York Academy of Sciences. https://doi.org/10.1111/nyas.13046

Fiebach, C. J., Vos, S. H., & Friederici, A. D. (2004). Neural correlates of syntactic ambiguity in sentence comprehension for low and high span readers. Journal of Cognitive Neuroscience, 16(9), 1562–1575.

Fischl, B., Rajendran, N., Busa, E., Augustinack, J., Hinds, O., Yeo, B. T. T., … Zilles, K. (2008). Cortical folding patterns and predicting cytoarchitecture. Cerebral Cortex. https://doi.org/10.1093/cercor/bhm225

Fitch, W. T., & Martins, M. D. (2014). Hierarchical processing in music, language, and action: Lashley revisited. Annals of the New York Academy of Sciences. https://doi.org/10.1111/nyas.12406

Frazier, L., & Rayner, K. (1987). Resolution of syntactic category ambiguities: Eye movements in parsing lexically ambiguous sentences. Journal of Memory and Language, 26(5), 505–526.

Friederici, A. D., Rüschemeyer, S.-A., Hahne, A., & Fiebach, C. J. (2003). The role of left inferior frontal and superior temporal cortex in sentence comprehension: localizing syntactic and semantic processes. Cerebral Cortex (New York, N.Y.: 1991), 13(2), 170–177. Retrieved from http://www.ncbi.nlm.nih.gov/pubmed/12507948

Friedrich, R., & Friederici, A. D. (2009). Mathematical logic in the human brain: Syntax. PLoS ONE. https://doi.org/10.1371/journal.pone.0005599

Frost, M. A., & Goebel, R. (2012). Measuring structural-functional correspondence: Spatial variability of specialised brain regions after macro-anatomical alignment. NeuroImage. https://doi.org/10.1016/j.neuroimage.2011.08.035

Futrell, R., Gibson, E., Tily, H. J. H., Blank, I., Vishnevetsky, A., Piantadosi, S. T., & Fedorenko, E. (2018). The Natural Stories Corpus. In Proceedings of the 11th Language Resources and Evaluation Conference (pp. 1–16). Miyazaki, Japan: European Language Resource Association. Retrieved from http://arxiv.org/abs/1708.05763

Gernsbacher, M. A. (1993). Less Skilled Readers Have Less Efficient Suppression Mechanisms. Psychological Science. https://doi.org/10.1111/j.1467-9280.1993.tb00567.x

Geschwind, N. (1970). The organization of language and the brain. Science. https://doi.org/10.1126/science.170.3961.940

Gibson, E. (1998). Linguistic complexity: locality of syntactic dependencies. Cognition, 68(1), 1–76. https://doi.org/10.1016/S0010-0277(98)00034-1

Gibson, E. (2000). The dependency locality theory: A distance-based theory of linguistic complexity. Image, Language, Brain, 95–126.

Gibson, E. A. F. (1991). A Computational Theory of Human Linguistic Processing: Memory Limitations and Processsing Breakdown, 206.

Gordon, P. C., Hendrick, R., & Levine, W. H. (2002). Memory-load interference in syntactic processing. Psychological Science. https://doi.org/10.1111/1467-9280.00475

Grodner, D., Gibson, E., & Tunstall, S. (2002). Syntactic Complexity in Ambiguity Resolution. Journal of Memory and Language, 46(2), 267–295. https://doi.org/10.1006/jmla.2001.2808

Hale, J. T., Lutz, D. E., Luh, W., Brennan, J. R., & Arbor, A. (2015). Modeling fMRI time courses with linguistic structure at various grain sizes. Proceedings of CMCL, 89–97.

Handwerker, D. A., Ollinger, J. M., & D’Esposito, M. (2004). Variation of BOLD hemodynamic responses across subjects and brain regions and their effects on statistical analyses. NeuroImage. https://doi.org/10.1016/j.neuroimage.2003.11.029

Hasson, U., Egidi, G., Marelli, M., & Willems, R. M. (2018). Grounding the neurobiology of language in first principles: The necessity of non-language-centric explanations for language comprehension. Cognition, 180, 135–157.

Hasson, U., & Honey, C. J. (2012). Future trends in {N}euroimaging: {N}eural processes as expressed within real-life contexts. NeuroImage, 62(2), 1272–1278.

Hasson, U., Yang, E., Vallines, I., Heeger, D. J., & Rubin, N. (2008). A hierarchy of temporal receptive windows in human cortex. Journal of Neuroscience, 28(10), 2539–2550.

Heim, S., Eickhoff, S. B., & Amunts, K. (2008). Specialisation in Broca’s region for semantic, phonological, and syntactic fluency? NeuroImage. https://doi.org/10.1016/j.neuroimage.2008.01.009

Henderson, J. M., Choi, W., Lowder, M. W., & Ferreira, F. (2016). Language structure in the brain: A fixation-related fMRI study of syntactic surprisal in reading. NeuroImage, 132, 293–300. https://doi.org/10.1016/j.neuroimage.2016.02.050

Henderson, J. M., Choi, W., Luke, S. G., & Desai, R. H. (2015). Neural correlates of fixation duration in natural reading: evidence from fixation-related fMRI. NeuroImage, 119, 390–397. https://doi.org/10.1016/j.neuroimage.2015.06.072

Holm, S. (1979). A simple sequentially rejective multiple test procedure. Scandinavian Journal of Statistics.

Hsu, A., Borst, A., & Theunissen, F. (2004). Quantifying variability in neural responses and its application for the validation of model predictions. Network: Computation in Neural Systems. https://doi.org/10.1088/0954-898x/15/2/002

Hsu, N. S., & Novick, J. M. (2016). Dynamic Engagement of Cognitive Control Modulates Recovery From Misinterpretation During Real-Time Language Processing. Psychological Science. https://doi.org/10.1177/0956797615625223

Humphries, C., Binder, J. R., Medler, D. A., & Liebenthal, E. (2006). Syntactic and semantic modulation of neural activity during auditory sentence comprehension. Journal of Cognitive Neuroscience. https://doi.org/10.1162/jocn.2006.18.4.665

Huth, A. G., de Heer, W. A., Griffiths, T. L., Theunissen, F. E., Gallant, J. L., Heer, W. a De, … Gallant, J. L. (2016). Natural speech reveals the semantic maps that tile human cerebral cortex. Nature, 532(7600), 453–458. https://doi.org/10.1038/nature17637.Natural

January, D., Trueswell, J. C., & Thompson-schill, S. L. (2009). NIH Public Access, 21(12), 2434–2444. https://doi.org/10.1162/jocn.2008.21179.Co-localization

January, D., Trueswell, J. C., & Thompson-Schill, S. L. (2009). Co-localization of Stroop and Syntactic Ambiguity Resolution in Broca’s Area: Implications for the Neural Basis of Sentence Processing. Journal of Cognitive Neuroscience, 21(12), 2434–2444. https://doi.org/10.1162/jocn.2008.21179

Johnson-Laird, P. N. (1983). Mental models: Towards a cognitive science of language, inference, and consciousness. Harvard University Press.

Julian, J. B., Fedorenko, E., Webster, J., & Kanwisher, N. (2012). An algorithmic method for functionally defining regions of interest in the ventral visual pathway. Neuroimage, 60(4), 2357–2364.

Jung-Beeman, M. (2005). Bilateral brain processes for comprehending natural language. Trends in Cognitive Sciences, 9(11), 512–518.

Just, M. A., & Carpenter, P. A. (1980). A theory of reading: From eye fixations to comprehension. Psychological Review. https://doi.org/10.1037/0033-295X.87.4.329

Just, M. a, Carpenter, P. a, & Woolley, J. D. (1982). Paradigms and processes in reading comprehension. Journal of Experimental Psychology. General, 111(2), 228–238. https://doi.org/10.1037/0096-3445.111.2.228

Kaakinen, J. K., & Hyönä, J. (2010). Task Effects on Eye Movements During Reading. Journal of Experimental Psychology: Learning Memory and Cognition. https://doi.org/10.1037/a0020693

Kaan, E., & Swaab, T. Y. (2002). The brain circuitry of syntactic comprehension. Trends in Cognitive Sciences. https://doi.org/10.1016/S1364-6613(02)01947-2

Keller, T. A., Carpenter, P. A., & Just, M. A. (2001). The Neural Bases of Sentence Comprehension: a fMRI Examination of Syntactic and Lexical Processing, 223–237.

Kennedy, A. (2000). Parafoveal processing in word recognition. Quarterly Journal of Experimental Psychology Section A: Human Experimental Psychology. https://doi.org/10.1080/713755901

King, J., & Just, M. A. (1991). Individual differences in syntactic processing: The role of working memory. Journal of Memory and Language, 30(5), 580–602.

Klein, R., & Farrell, M. (1989). Search performance without eye movements. Perception & Psychophysics. https://doi.org/10.3758/BF03210863

Kuperberg, G. R., Holcomb, P. J., Sitnikova, T., Greve, D., Dale, A. M., & Caplan, D. (2003). Distinct patterns of neural modulation during the processing of conceptual and syntactic anomalies. Journal of Cognitive Neuroscience, 15(2), 272–293. https://doi.org/10.1162/089892903321208204

Kuperberg, G. R., Sitnikova, T., & Lakshmanan, B. M. (2008). Neuroanatomical distinctions within the semantic system during sentence comprehension: evidence from functional magnetic resonance imaging. Neuroimage, 40(1), 367–388.

Lerner, Y., Honey, C. J., Silbert, L. J., & Hasson, U. (2011). Topographic mapping of a hierarchy of temporal receptive windows using a narrated story. The Journal of Neuroscience, 31(8), 2906–2915.

Lescroart, M. D., & Gallant, J. L. (2019). Human Scene-Selective Areas Represent 3D Configurations of Surfaces. Neuron. https://doi.org/10.1016/j.neuron.2018.11.004

Lescroart, M. D., Stansbury, D. E., & Gallant, J. L. (2015). Fourier power, subjective distance, and object categories all provide plausible models of BOLD responses in scene-selective visual areas. Frontiers in Computational Neuroscience, 9, 135. https://doi.org/10.3389/fncom.2015.00135

Levy, R. (2008). Expectation-based syntactic comprehension. Cognition, 106(3), 1126–1177. https://doi.org/10.1016/j.cognition.2007.05.006

Lewis, R. L., & Vasishth, S. (2005). An activation-based model of sentence processing as skilled memory retrieval. Cognitive Science. https://doi.org/10.1207/s15516709cog0000_25

Lewis, R. L., Vasishth, S., & Van Dyke, J. A. (2006). Computational principles of working memory in sentence comprehension. Trends in Cognitive Sciences, 10(10), 447–454. https://doi.org/10.1016/j.tics.2006.08.007

Lopopolo, A., Frank, S. L., Van Den Bosch, A., & Willems, R. M. (2017). Using stochastic language models (SLM) to map lexical, syntactic, and phonological information processing in the brain. PLoS ONE, 12(5), 1–18. https://doi.org/10.1371/journal.pone.0177794

Mahowald, K., & Fedorenko, E. (2016). Reliable individual-level neural markers of high-level language processing: A necessary precursor for relating neural variability to behavioral and genetic variability. NeuroImage. https://doi.org/10.1016/j.neuroimage.2016.05.073

Mazoyer, B. M., Tzourio, N., Frak, V., Syrota, A., Murayama, N., Levrier, O., … Mehler, J. (1993). The cortical representation of speech. Journal of Cognitive Neuroscience, 5(4), 467–479. https://doi.org/10.1162/jocn.1993.5.4.467

McElree, B. (2000). Sentence comprehension is mediated by content-addressable memory structures. Journal of Psycholinguistic Research. https://doi.org/10.1023/A:1005184709695

McElree, B. (2001). Working Memory and Focal Attention. Journal of Experimental Psychology: Learning Memory and Cognition. https://doi.org/10.1037/0278-7393.27.3.817

McMillan, C. T., Clark, R., Gunawardena, D., Ryant, N., & Grossman, M. (2012). FMRI evidence for strategic decision-making during resolution of pronoun reference. Neuropsychologia. https://doi.org/10.1016/j.neuropsychologia.2012.01.004

McMillan, C. T., Coleman, D., Clark, R., Liang, T. W., Gross, R. G., & Grossman, M. (2013). Converging evidence for the processing costs associated with ambiguous quantifier comprehension. Frontiers in Psychology, 4(APR), 1–10. https://doi.org/10.3389/fpsyg.2013.00153

Menke, J., & Martinez, T. R. (2004). Using permutations instead of student’s t distribution for p-values in paired-difference algorithm comparisons. In 2004 IEEE International Joint Conference on Neural Networks (IEEE Cat. No. 04CH37541) (Vol. 2, pp. 1331–1335).

Mesulam, M. M. (1998). From sensation to cognition. Brain. https://doi.org/10.1093/brain/121.6.1013

Miller, E. K., & Cohen, J. D. (2001). An Integrative Theory of Prefrontal Cortex Function. Annual Review of Neuroscience. https://doi.org/10.1146/annurev.neuro.24.1.167

Mineroff, Z., Blank, I. A., Mahowald, K., & Fedorenko, E. (2018). A robust dissociation among the language, multiple demand, and default mode networks: Evidence from inter-region correlations in effect size. Neuropsychologia, 119, 501–511.

Mitchell, D. C. (1984). An evaluation of subject-paced reading tasks and other methods for investigating immediate processes in reading. New Methods in Reading Comprehension Research, 69–89.

Mollica, F., Siegelman, M., Diachek, E., Piantadosi, S. T., Mineroff, Z., Futrell, R., … Fedorenko, E. (2020). Composition is the core driver of the language-selective network. Neurobiology of Language. https://doi.org/10.1162/nol_a_00005

Murphy, B., Hale, J., & Brennan, J. (2016). Grammatical Relations in the Listening Brain. Poster at PRNI 2016, Pattern Recognition and Neuroimaging Conference.

Naselaris, T., Kay, K. N., Nishimoto, S., & Gallant, J. L. (2011). Encoding and decoding in fMRI. NeuroImage, 56(2), 400–410. https://doi.org/10.1016/j.neuroimage.2010.07.073

Nelson, M. J., El Karoui, I., Giber, K., Yang, X., Cohen, L., Koopman, H., … Dehaene, S. (2017). Neurophysiological dynamics of phrase-structure building during sentence processing. Proceedings of the National Academy of Sciences of the United States of America. https://doi.org/10.1073/pnas.1701590114

Nieto-Castañón, A., & Fedorenko, E. (2012). Subject-specific functional localizers increase sensitivity and functional resolution of multi-subject analyses. NeuroImage. https://doi.org/10.1016/j.neuroimage.2012.06.065

Nieuwland, M. S., Martin, A. E., & Carreiras, M. (2012). Brain regions that process case: Evidence from basque. Human Brain Mapping, 33(11), 2509–2520. https://doi.org/10.1002/hbm.21377

Novais-Santos, S., Gee, J., Shah, M., Troiani, V., Work, M., & Grossman, M. (2007). Resolving sentence ambiguity with planning and working memory resources: Evidence from fMRI. NeuroImage, 37(1), 361–378. https://doi.org/10.1016/j.neuroimage.2007.03.077

Novick, J. M., Kan, I. P., Trueswell, J. C., & Thompson-Schill, S. L. (2009). A case for conflict across multiple domains: Memory and language impairments following damage to ventrolateral prefrontal cortex. Cognitive Neuropsychology. https://doi.org/10.1080/02643290903519367

Novick, J. M., Trueswell, J. C., & Thompson-Schill, S. L. (2005). Cognitive control and parsing: Reexamining the role of Broca’s area in sentence comprehension. Cognitive, Affective and Behavioral Neuroscience. https://doi.org/10.3758/CABN.5.3.263

Pallier, C., Devauchelle, A. D., & Dehaene, S. (2011). Cortical representation of the constituent structure of sentences. Proceedings of the National Academy of Sciences, 108(6), 2522–2527.

Patel, A. D. (2003). Language, music, syntax and the brain. Nature Neuroscience. https://doi.org/10.1038/nn1082

Patel, A. D. (2012). Music, Language, and the Brain. Music, Language, and the Brain. https://doi.org/10.1093/acprof:oso/9780195123753.001.0001

Paunov, A. M., Blank, I. A., & Fedorenko, E. (2019). Functionally distinct language and Theory of Mind networks are synchronized at rest and during language comprehension. Journal of Neurophysiology, 121(4), 1244–1265.

Peelle, J. E., Troiani, V., Wingfield, A., & Grossman, M. (2010). Neural processing during older adults’ comprehension of spoken sentences: Age differences in resource allocation and connectivity. Cerebral Cortex, 20(4), 773–782. https://doi.org/10.1093/cercor/bhp142

Posner, M. I. (1980). Orienting of attention. The Quarterly Journal of Experimental Psychology. https://doi.org/10.1080/00335558008248231

Posner, Michael I. (2016). Orienting of attention: Then and now. Quarterly Journal of Experimental Psychology. https://doi.org/10.1080/17470218.2014.937446

Rasmussen, N. E., & Schuler, W. (2018). Left-Corner Parsing With Distributed Associative Memory Produces Surprisal and Locality Effects. Cognitive Science. https://doi.org/10.1111/cogs.12511

Rayner, K. (1977). Visual attention in reading: Eye movements reflect cognitive processes. Memory & Cognition. https://doi.org/10.3758/BF03197383

Rayner, K. (1978). Eye movements in reading and information processing. Psychological Bulletin. https://doi.org/10.1037/0033-2909.85.3.618

Rayner, K. (1998). Eye movements in reading and information processing: 20 years of research. Psychological Bulletin, 124(3), 372. https://doi.org/10.1037/0033-2909.124.3.372

Remington, R. W. (1980). Attention and saccadic eye movements. Journal of Experimental Psychology: Human Perception and Performance. https://doi.org/10.1037/0096-1523.6.4.726

Resnik, P. (1992). Left-corner parsing and psychological plausibility. https://doi.org/10.3115/992066.992098

Rodd, J. M., Davis, M. H., & Johnsrude, I. S. (2005). The neural mechanisms of speech comprehension: fMRI studies of semantic ambiguity. Cerebral Cortex, 15(8), 1261–1269. https://doi.org/10.1093/cercor/bhi009

Rodd, J. M., Johnsrude, I. S., & Davis, M. H. (2010). The role of domain-general frontal systems in language comprehension: Evidence from dual-task interference and semantic ambiguity. Brain and Language. https://doi.org/10.1016/j.bandl.2010.07.005

Rodriguez, A., & Granger, R. (2016). The grammar of mammalian brain capacity. Theoretical Computer Science. https://doi.org/10.1016/j.tcs.2016.03.021

Rogalsky, C., & Hickok, G. (2011). The role of Broca’s area in sentence comprehension. Journal of Cognitive Neuroscience. https://doi.org/10.1162/jocn.2010.21530

Saxe, R., Brett, M., & Kanwisher, N. (2006). Divide and conquer: A defense of functional localizers. NeuroImage. https://doi.org/10.1016/j.neuroimage.2005.12.062

Schotter, E. R., Tran, R., & Rayner, K. (2014). Don’t believe what you read (Only Once): Comprehension is supported by regressions during reading. Psychological Science. https://doi.org/10.1177/0956797614531148

Schuler, W., AbdelRahman, S., Miller, T., & Schwartz, L. (2010). Broad-coverage parsing using human-like memory constraints. Computational Linguistics. https://doi.org/10.1162/coli.2010.36.1.36100

Scott, T. L., Gallée, J., & Fedorenko, E. (2017). A new fun and robust version of an fMRI localizer for the frontotemporal language system. Cognitive Neuroscience, 8(3), 167–176.

Shain, C., Blank, I. A., van Schijndel, M., Schuler, W., & Fedorenko, E. (2019). fMRI reveals language-specific predictive coding during naturalistic sentence comprehension. BioRxiv, 717512. https://doi.org/10.1016/j.neuropsychologia.2019.107307

Shain, C., Blank, I. A., van Schijndel, M., Schuler, W., & Fedorenko, E. (2020). fMRI reveals language-specific predictive coding during naturalistic sentence comprehension. Neuropsychologia. https://doi.org/10.1016/j.neuropsychologia.2019.107307

Slevc, L. R., Rosenberg, J. C., & Patel, A. D. (2009). Making psycholinguistics musical: Self-paced reading time evidence for shared processing of linguistic and musical syntax. Psychonomic Bulletin and Review. https://doi.org/10.3758/16.2.374

Smith, N. J., & Levy, R. (2013). The effect of word predictability on reading time is logarithmic. Cognition, 128(3), 302–319. https://doi.org/10.1016/j.cognition.2013.02.013

Snijders, T. M., Vosse, T., Kempen, G., Van Berkum, J. J. A., Petersson, K. M., & Hagoort, P. (2009). Retrieval and unification of syntactic structure in sentence comprehension: An fMRI study using word-category ambiguity. Cerebral Cortex, 19(7), 1493–1503. https://doi.org/10.1093/cercor/bhn187

Speer, N K, Zacks, J. M., & Reynolds, J. R. (2007). Human brain activity time-locked to narrative even boundaries. Psychol. Sci., 18(5), 449–455. Retrieved from http://dx.doi.org/10.1111/j.1467-9280.2007.01920.x

Speer, Nicole K, Reynolds, J. R., Swallow, K. M., & Zacks, J. M. (2009). Reading stories activates neural representations of visual and motor experiences. Psychological Science, 20(8), 989–999. https://doi.org/10.1111/j.1467-9280.2009.02397.x.Reading

Sreenivasan, K. K., Curtis, C. E., & D’Esposito, M. (2014). Revisiting the role of persistent neural activity during working memory. Trends in Cognitive Sciences. https://doi.org/10.1016/j.tics.2013.12.001

Stowe, L. A., Broere, C. A. J., Paans, A. M. J., Wijers, A. A., Mulder, G., Vaalburg, W., & Zwarts, F. (1998). Localizing components of a complex task: Sentence processing and working memory. NeuroReport. https://doi.org/10.1097/00001756-199809140-00014

Tahmasebi, A. M., Davis, M. H., Wild, C. J., Rodd, J. M., Hakyemez, H., Abolmaesumi, P., & Johnsrude, I. S. (2012). Is the link between anatomical structure and function equally strong at all cognitive levels of processing? Cerebral Cortex. https://doi.org/10.1093/cercor/bhr205

Taylor, J. S. H., Rastle, K., & Davis, M. H. (2014). Interpreting response time effects in functional imaging studies. NeuroImage. https://doi.org/10.1016/j.neuroimage.2014.05.073

Tettamanti, M., & Weniger, D. (2006). Broca’s area: A supramodal hierarchical processor? Cortex. https://doi.org/10.1016/S0010-9452(08)70384-8

Thesen, S., Heid, O., Mueller, E., & Schad, L. R. (2000). Prospective Acquisition Correction for head motion with image-based tracking for real-time fMRI. Magnetic Resonance in Medicine, 44(3), 457–465. https://doihttps://doi.org/10.1002/1522-2594(200009)44:3<457::AID-MRM17>3.0.CO;2-R

Thompson-Schill, S. L., Bedny, M., & Goldberg, R. F. (2005). The frontal lobes and the regulation of mental activity. Current Opinion in Neurobiology. https://doi.org/10.1016/j.conb.2005.03.006

Vagharchakian, L., Dehaene-Lambertz, G., Pallier, C., & Dehaene, S. (2012). A temporal bottleneck in the language comprehension network. Journal of Neuroscience, 32(26), 9089–9102.

van Schijndel, M., Exley, A., & Schuler, W. (2013). A Model of Language Processing as Hierarchic Sequential Prediction. Topics in Cognitive Science. https://doi.org/10.1111/tops.12034

Vandenberghe, R., Nobre, A. C., & Price, C. J. (2002). The response of left temporal cortex to sentences. Journal of Cognitive Neuroscience. https://doi.org/10.1162/08989290260045800

Vasishth, S., von der Malsburg, T., & Engelmann, F. (2013). What eye movements can tell us about sentence comprehension. Wiley Interdisciplinary Reviews: Cognitive Science, 4(2), 125–134.

Vázquez-Rodríguez, B., Suárez, L. E., Markello, R. D., Shafiei, G., Paquola, C., Hagmann, P., … Misic, B. (2019). Gradients of structure–function tethering across neocortex. Proceedings of the National Academy of Sciences of the United States of America. https://doi.org/10.1073/pnas.1903403116

Vergauwe, E., Barrouillet, P., & Camos, V. (2010). Do mental processes share a domain-general resource? Psychological Science. https://doi.org/10.1177/0956797610361340

Waters, G. S., & Caplan, D. (1996). The Measurement of Verbal Working Memory Capacity and Its Relation to Reading Comprehension. Quarterly Journal of Experimental Psychology Section A: Human Experimental Psychology. https://doi.org/10.1080/713755607

Wehbe, L., Murphy, B., Talukdar, P., Fyshe, A., Ramdas, A., & Mitchell, T. (2014). Simultaneously uncovering the patterns of brain regions involved in different story reading Subprocesses. PloS One, 9(11), e112575. https://doi.org/10.1371/journal.pone.0112575

Weichwald, S., Meyer, T., Özdenizci, O., Schölkopf, B., Ball, T., & Grosse-Wentrup, M. (2015). Causal interpretation rules for encoding and decoding models in neuroimaging. Neuroimage, 110, 48–59.

Wernicke, C. (1874). Der aphasische Symptomencomplex. Eine psychologische Studie auf anatomischer Basis. [The aphasia symptom complex. A psychological study on an anatomical basis]. Wernicke’s work on aphasia.

Whitfield-Gabrieli, S., & Nieto-Castanon, A. (2012). Conn: a functional connectivity toolbox for correlated and anticorrelated brain networks. Brain Connectivity, 2(3), 125–141.

Whitney, C., Huber, W., Klann, J., Weis, S., Krach, S., & Kircher, T. (2009). Neural correlates of narrative shifts during auditory story comprehension. NeuroImage, 47(1), 360–366. https://doi.org/10.1016/j.neuroimage.2009.04.037

Wild, C. J., Yusuf, A., Wilson, D. E., Peelle, J. E., Davis, M. H., & Johnsrude, I. S. (2012). Effortful listening: The processing of degraded speech depends critically on attention. Journal of Neuroscience. https://doi.org/10.1523/JNEUROSCI.1528-12.2012

Willems, R. M., Frank, S. L., Nijhof, A. D., Hagoort, P., & Van Den Bosch, A. (2016). Prediction during Natural Language Comprehension. Cerebral Cortex, 26(6), 2506–2516. https://doi.org/10.1093/cercor/bhv075

Wright, P., Randall, B., Marslen-Wilson, W. D., & Tyler, L. K. (2011). Dissociating linguistic and task-related activity in the left inferior frontal gyrus. Journal of Cognitive Neuroscience, 23(2), 404–413.

Wright, R., & Ward, L.. (2008). Eye movements and attention shifts. Orienting of attention.

Yarkoni, T., Barch, D. M., Gray, J. R., Conturo, T. E., & Braver, T. S. (2009). BOLD correlates of trial-by-trial reaction time variability in gray and white matter: A multi-study fMRI analysis. PLoS ONE, 4(1). https://doi.org/10.1371/journal.pone.0004257

